# Imaging spectroscopy reveals topographic variability effects on grassland functional traits and drought responses

**DOI:** 10.1101/2023.12.31.573803

**Authors:** Phuong D. Dao, Yuhong He, Bing Lu, Alexander Axiotis

**Author notes:** Author for correspondence: Phuong D. Dao.

## Abstract

Functional traits and their variations are essential indicators of plant metabolism, growth, distribution, and survival and determine how a plant and an ecosystem function. Under the same climatic condition, traits can vary largely between species and within the same species growing in different topographic conditions. When drought stress occurs, plants that grow in these conditions may respond differently as their topography-driven tolerance and adaptability differ. Insights into topographic variability-driven trait variation and drought response can improve our prediction of ecosystem functioning and ecological impacts. Imaging spectroscopy allows accurate detection of plant species, retrieval of functional traits, and characterization of topography-driven and drought impacts on trait variation across space. However, the use of this data in a heterogeneous grassland ecosystem is challenging as species are small, high mixed, spectrally and texturally similar, and highly varied with small-scale variation in topography. In this paper, we introduce the first study that explores the use of high-resolution airborne imaging spectroscopy to characterize the variation of common traits, including chlorophylls (Chl), carotenoids (Car), Chl/Car ratio, water content (WC), and leaf area index (LAI), across topographic gradients and under drought stress at the species level in a heterogeneous grassland. The results reveal that there were significant relationships between functional traits and topographic variability, and the degree of the relationships deferred among species and under different environmental conditions. The results also show that drought-induced trait responses varied significantly within and between species, especially between drought-tolerant invasive species and native species, between lower and upper slope positions. The study contributes greatly to the advancement in understanding biological and ecological processes for a better prediction of ecosystem functioning under stressed environments.

## 1 INTRODUCTION

Functional traits (e.g., pigments, water content, specific leaf area, nitrogen, non-structural carbohydrates) play essential roles in plant metabolism, growth, distribution, and survival (Roscher et al. 2012, Asner et al. 2017, Maharjan et al. 2021). Variations in functional traits reflect how plants adapt to environmental change and respond to biotic and abiotic disturbances, including drought stress (Trlica and Biondini 1990, Dong et al. 2014, Asner et al. 2017). Droughts pose serious threats to plants by inhibiting photosynthesis, causing cell damages, reducing productivity, and triggering mortality (Monclus et al. 2006, McDowell et al. 2008). In response, plants with special adaptive traits can develop strategies to protect themselves from damages. For example, leaf stomata close to reduce water loss (Taiz and Zeiger 2002). Xanthophyll-cycle pigment pool size increases rapidly, and violaxanthin is de-epoxidized to zeaxanthin, an energy-quenching pigment, to help dissipate excess radiation to protect photosynthetic apparatus from oxidative damages (Demmig-Adams 1990, Jahns and Holzwarth 2012). Leaves can then roll to reduce leaf area to minimize radiation load (Trlica and Biondini 1990). However, drought response is species-specific, depending on their tolerance and plasticity, especially between native and invasive species (Bueno et al. 2020). Some invasive species may develop unique strategies and traits to adapt to resist droughts. It is observed that when these plants invade a dry environment, they quickly adjust to the living condition, and outcompete, suppress, and replace native plants (Dyer and Rice 1999, Trlica and Biondini 1990, Gaskin et al. 2020). Trlica and Biondini (1990) discovered that leaves of invasive *Agropyron cristatum* roll more rapidly than native *Agropyron smithii* and *Bouteloua gracilis* to reduce radiation load when experienced drought stress. Invasive smooth brome has higher root-to-shoot allocation than native *Pascopyrum smithii* and *Stipa viridula Trin.* Under drought stress, indicating its higher drought-resistant capacity (Dong et al. 2014). Hence, insights about trait-related drought response of invasive and native species help us better understand biological and ecological processes for improved prediction of plant functioning, plant growth, and production under stressed environments.

There is a strong relationship between topography and soil conditions such as soil water runoff, soil moisture, nutrient richness (Bohlman et al. 2008, Homeier et al. 2010), especially in rugged terrain due to heterogeneous microclimate and soil hydrologic and edaphic conditions (Pierick et al. 2020, Takyu et al. 2003). For example, soil at low slope positions often has higher moisture, richer nutrient, and lower water runoff rates than soil at upper slope positions (Balvanera et al. 2011, Gibbons and Newbery 2003). This topographic variability can have significant effects on the variation of traits such as pigment content, leaf size, leaf area, leaf mass per area, and leaf thickness (Ackerly et al. 2002, Cornwell and Ackerly 2009, Schmitt et al. 2020), and root traits including root diameter, length, and tissue density (Pierick et al. 2020). Such effects subsequently control plant performance, growth, productivity, and spatial distribution pattern (Homeier et al. 2010, Adams et al. 2014, Bohlman et al. 2008, Girardin et al. 2014). Moreover, plants over upper slope positions with limited water availability and nutrients may adjust their traits to adapt to withstand water and nutrient scarcity compared to those at low slope regions. This trait adjustment and adaptation may lead to distinctions in within-species and between-species drought response, especially between native and invasive species. However, differences in topography-driven drought response among species have not been well-documented in the literature (Schmitt et al. 2020). This knowledge gap limits our ability to understand plant and ecosystem processes and functions under drought conditions. Furthermore, the majority of previous studies that relied on laboratory and plot measurements (Pierick et al. 2020, Schmitt et al. 2020, Pfennigwerth et al. 2017, Ackerly et al. 2002) were unable to capture the trait variation over space. Not only that, plot-level averaged measurements would undermine trait variation in grassland ecosystems as a plot may comprise multiple small and co-existing species.

Imaging spectroscopy has been used to retrieve plant biophysical, biochemical, and structural characteristics (Lu et al. 2019, Serbin and Townsend 2020) and assess drought-induced changes (Wen et al. 2021, Carter and Knapp 2001) across various spatial and temporal scales. Particularly, airborne imaging spectroscopy with high spatial resolutions and hundreds to thousands of narrow spectral bands is able to distinguish subtle changes in target objects (Dao et al. 2021b, Lu et al. 2020). As optical properties of vegetation are controlled by leaf structure, water content, and biochemical and canopy structural properties at specific wavelength regions (Asner 1998, Gates et al. 1965), drought-induced changes in these traits can be detected from spectral response. These traits can be mapped through empirical Ordinary Least Squares (OLS) regression against spectral indices (Gilerson et al. 2010) or through Partial Least Squares Regression (PLSR, (Wold et al. 2001)) against multiple spectral indices (Lu et al. 2019) or full continuous spectra (Singh et al. 2015, Serbin et al. 2014). Trait maps derived from remote sensing facilitate the assessment of trait variation within and between species, especially between native and invasive species, and drought-induced changes across space.

Several studies have investigated the impact of topography on foliar traits using imaging spectroscopy in combination with a digital elevation model (DEM). A recent study by Asner et al. (2015) explored the effect of topography and soil types on plant traits at a grain size of 1 ha. A further study (Asner et al. 2017) evaluated the spatial-scale dependency of traits at the grain sizes of 0.01 ha and 1 ha. Another study by Swinfield et al. (2020) examined the effect of topography and logging on foliar traits of trees in tropical forests. However, the majority of these studies either focused on forests or were conducted over large geographic regions that encompass various soil types and soil fertility, climate gradients, plant functional types, or even biomes. These large-scale analyses can provide overall information about stress-induced variation in a plant community or an ecosystem but would underestimate trait variation in highly mixed grassland ecosystems, of which species composition varies at centimeter or decimeter scales (Lu and He 2018, Dao et al. 2021a). This is due to the mixed signals and generalized measurements of multiple species and topography-driven trait variation within a coarse resolution pixel. Asner et al. (2017) also showed the dependence of foliar traits on sampling scale along topographic gradients. The generalized measurements in previous studies would underestimate the underlying trait variation and drought response within and between species driven by small-scale topographic variability. This would limit our ability to accurately predict ecosystem processes and functions in stressed environments. To the best of our knowledge, no studies have investigated how and to what extent small-scale topographic variability affects trait variation and drought-induced trait response at species level, especially between native and invasive grassland species, with high-resolution imagery.

In this study, we integrated airborne high-resolution imaging spectroscopy and DEM data to explore how small-scale variability in topography determines within-species and between-species trait variation and drives the drought-induced trait response in native and invasive grassland species. This study aims to answer three following research questions:

1. To what extent, can leaf and canopy functional traits of native and invasive species in a highly mixed grassland ecosystem be quantitatively retrieved from leaf reflectance and airborne imaging spectroscopy?
2. How does small-scale topographic variability affect within-species and between-species trait variation under normal and drought conditions across topographic gradients and classified categories?
3. How do drought trait responses differ within and between species, especially between native and drought-tolerant invasive species, across topographic gradients and classified categories?

## 2 MATERIALS AND METHODS

### 2.1 Study site

The study was conducted at Koffler Scientific Reserve in Ontario, Canada (Figure 1). The area is dominated by two invasive species including smooth brome (*Bromus inermis*) and orchard grass (*Dactylis glomerata*) and three native species, including goldenrod (*Solidago canadensis*), common milkweed (*Asclepias syriaca*), and wild grape (*Vitis riparia*). These species are perennial plants, and their distribution and composition are the same over time (e.g., 2016 to 2017 when the surveys were conducted). Our field vegetation survey also confirms the patterns. Fine sandy loams and silty sand loams with the accumulation of organic matter in lowland regions are the major soil types in the area. The hilly opography varies largely from low slope and elevation (279 m) to upper slope and high elevation (310 m). The site belongs to the warm-summer humid continental climate with hot and humid summers and cold winters (Peel et al. 2007, Beck et al. 2018). The climate normal data at King Smoke Tree weather station (about 2 km from the reserve) show the daily average temperature is about-7.4°C in January and 20.5°C in July, and the average precipitation is approximately 850 mm per year.

**Figure 1.**
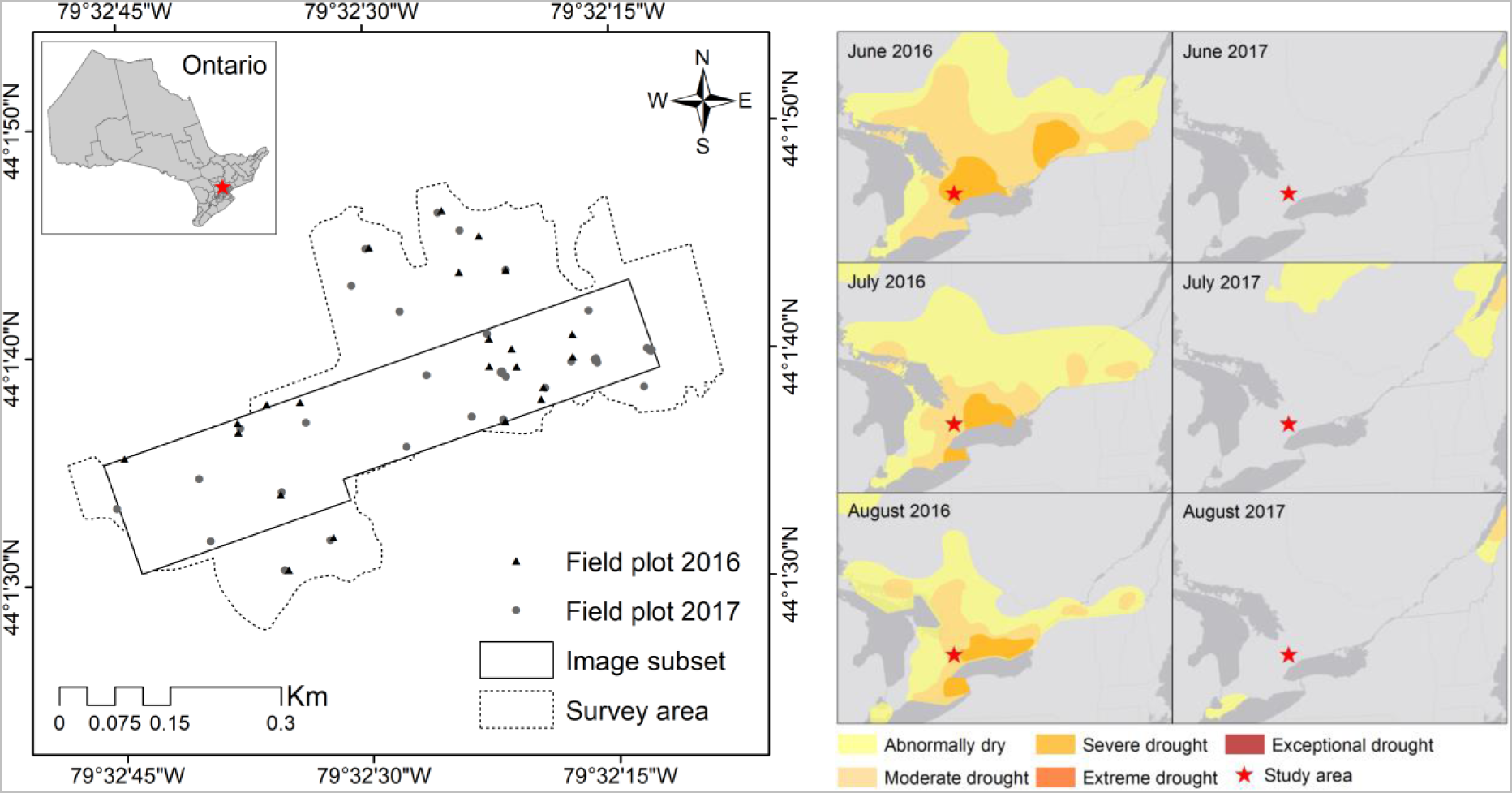
The study area (left figure) in Koffler Scientific Reserve, Ontario, with locations of field vegetation survey plots. The dashed line is the boundary of the field surveyed grassland area, the solid line is the boundary of the area of airborne image acquisition, black triangles and gray circles are field survey plots of 2016 and 2017, respectively. Right figures are maps of drought severity in June, July, and August of 2016 and 2017 in and around the study area provided by the Canadian Drought Monitor program, Agriculture and Agri-Food Canada.

Continuous drought monitoring data from the Canadian Drought Monitoring program, Agriculture and Agri-Food Canada (https://agriculture.canada.ca/en/canadian-drought-monitor) for 2016 and 2017, were shown in Figure 1. The drought maps were derived from the combination of precipitation, temperature, Standardized Precipitation Index, Palmer Drought Index, streamflow data, satellite-derived vegetation index, and other climate data (Tadesse et al. 2017, Agriculture and Agri-Food Canada 2016). The maps show the area was affected by a severe drought in June and moderate drought in July and August 2016, while there was no drought in 2017. Besides, ground-measured soil moistures at most of the field plots were lower than 18%, with many values being less than 10% in 2016. These levels of soil moisture can cause a significant reduction in aboveground biomass and plant nitrogen concentration in grasses and forbs (Mackie et al. 2019). While the moisture of most plots was higher than 20%, only a few values from high topographic positions were below 20% in 2017. Our multiyear surveys and measurements show that drought responses within each species and between species varied across topographic gradients. As the climatic condition was the same across space in this small area while topography varies greatly, we hypothesize that small-scale variation in topography was the major factor that drove the within-and between-species trait variation and controlled drought-induced trait response in these species.

### 2.2 Plant trait measurements

In this study, six common traits, including five leaf-level traits (Chl, Car, Chl/Car, WC, and specific leaf area–SLA) and five canopy-level traits (Chl, Car, Chl/Car, WC, and LAI) were retrieved from leaf reflectance and airborne images, respectively. It should be noted that the implementation of the leaf-level trait retrieval was only for confirming the utility of leaf reflectance in quantifying traits, not for topography-trait relationship assessment. Only canopy-level traits were used to link with the topographic variable to assess the topographic effect since this study primarily focuses on evaluating the performance of airborne imagery. During the field surveys, LAIs of four corners and the centre of 3×3 m plots (22 plots in 2016 and 40 plots in 2017) were recorded on the same day of airborne imaging (23 August 2016 and 20 August 2017) with an AccuPAR Ceptometer from Decagon Devices, Inc., Pullman, Washington, USA. Foliar samples were collected at the same field plots and on the same days of airborne imaging for leaf trait measurements. At each plot, leaf samples of different greenness and heights of each of the five species were collected. In total, 244 leaf samples (102 samples for the 2016 campaign and 142 samples for the 2017 campaign) were collected. A portion of each leaf was dried up in an oven under 80°C for 24 hours for leaf WC (WC = (fresh weight-dry weight)/sample area) and SLA (SLA = sample area/dry mass) calculation. We put another portion of the leaf with an area of 0.7694 cm^2^ (clipped using a hole puncher) in 4 ml of Dimethylformamide solvent under-4°C for 2 days and then measured the absorbance of the dissolved liquid at 480 nm, 646.8 nm, and 663.8 nm (Dao et al. 2021a) using a GENESYS 10S UV-Vis spectrophotometer from Thermo Fisher Scientific Inc., Wisconsin, USA. Total Chl and Car were then calculated from the absorbance using equations developed by (Porra et al. 1989, Wellburn 1994). Leaf Chl, Car, and WC were then scaled up to the canopy level, to be linked with airborne images, by multiplying with the corresponding LAI values.

### 2.3 Field and leaf measurements

Ground reflectance measurements of reference materials as described in (Dao et al. 2019a), were collected during image acquisition using a FieldSpec 3 spectroradiometer (Malvern PANalytical Company, United Kingdom) for radiometric correction of images. The sensor measured reflectance from 400 nm to 2500 nm at an interval of 1 nm. A Spectralon white reference was used for calibration. For each reference target, 15 reflectance measurements were collected at four corners and the centre. The reflectances were resampled to 301 bands ranging from 400 nm to 1000 nm with a spectral interval of 2 nm to match with the spectral resolution of airborne images.

During the imaging campaigns, 29 ground control points (GCPs) were also collected using the real-time kinematic Global Navigation Satellite System technique (RTK-GNSS, with two single-frequency (L1) EMLID Reach RS^+^ receivers) for image registration. The RTK technique provided GCPs with a post-processing horizontal accuracy of 0.15–0.25 m.

In addition to the use of airborne imaging spectroscopy in canopy-level traits, leaf-level reflectances were also used to confirm the potential of retrieving foliar traits at the leaf level. Leaf reflectances were measured on 244 leaf samples, that were later used for pigments and water measurements, using the FieldSpec 3. The sensor was equipped with a leaf clip that produces consistent full-spectrum halogen light and has a Spectralon white reference panel for calibration. For each sample, we recorded three measurements, resulting in 732 reflectances recorded in total.

### 2.4 Airborne image acquisition and preprocessing

Hyperspectral images were acquired on 23 August 2016 and 20 August 2017 by a Mirco-Hyperspec VNIR push-broom sensor (Headwall Photonics Inc, USA), operated on a JetRanger helicopter. The sensor has 325 narrow bands ranging from 400 nm to 1000 nm, with a spectral resolution of about 2 nm and a radiometric resolution of 12 bits. The imaging system was also equipped with a global positioning system (GPS) and an inertial measurement unit (IMU) that recorded the sensor’s coordinates, sensor’s orientation (e.g., roll, pitch, yaw), and altitudes for geo-correction. The sensor was operated at about 200 m above the ground, capturing images at a spatial resolution of about 0.2 m. Images were captured under stable sunlight conditions without cloud cover–between 10:30-11:00 am local time.

Raw images were first geometrically corrected with a built-in DEM, and digital numbers were then converted to at-sensor radiance in the SpectralView software (Headwall Photonics Inc, USA). Images were then projected to the Universal Transverse Mercator (UTM) system, Zone 17N in the North American Datum 1983 (NAD83). Images were registered using GCPs and mosaicked to create the complete image of the study area. Both ground reference reflectance and hyperspectral images were resampled to 301 bands (400-1000 nm) with a spectral interval of 2 nm. Finally, images were radiometrically corrected using the empirical line calibration procedure proposed by Dao et al. (2019a) and smoothed using the Savitzky-Golay filter (Savitzky and Golay 1964).

### 2.5 Species classification

To characterize trait variation within each species and between species, mapping species distribution and composition is an essential step. From multiyear inventory, we observed that the distribution and composition of the five perennial species remained stable at a given time between the two years (23 August 2016 and 20 August 2017). Hence, species classification was done on the 2017 image as species were healthy at their full canopy and had underlying spectral distinctions, resulting in accurate classification results. The species classification was implemented using an object-based image analysis (OBIA) method (Blaschke 2010, Dao et al. 2019b) that consists of an image segmentation step using the compact watershed algorithm (Beucher and Lantuéjoul 1979) with noise normalization (Dao et al. 2021b) and an image classification step using a Random Forest (RF) classifier (Breiman 2001). The algorithm was implemented on the three principal component analysis (PCA) transformed components, which explained 99.9% of the variance of the original image. The watershed segmentation was implemented using the Scikit-image library (Van der Walt et al. 2014), and the RF classification was implemented using the Scikit-learn library (Pedregosa et al. 2011) in Python. The model input was the combination of the first-order derivative spectra of the hyperspectral image (Dao et al. 2021a) and six textural layers, including correlation, contrast, dissimilarity, entropy, homogeneity, and variance (Farwell et al. 2020). The classification model was trained with a training set, and the model’s optimal parameters were determined through a random search with k-fold cross-validation (k=10). A total number of 350 ground reference samples (collected using RTK-GNSS) were randomly split into a training set (80%) for model training and a test set (20%) for accuracy assessment. The classification results were evaluated through four commonly used metrics, including Producer’s Accuracy (PA), User’s Accuracy (UA), Overall Accuracy (OA), and Cohen’s Kappa coefficient (Kappa) (Cohen 1960, Congalton and Green 2008). Details on model training, image classification, and accuracy assessment were discussed in Dao et al. (2021a).

### 2.6 Plant trait retrieval from spectra

Leaf-and canopy-level traits were retrieved using kernel Partial Least Squares Regression (Wold et al. 2001, Dayal and MacGregor 1997) that links leaf reflectance and airborne image reflectance with each of the plant traits. PLSR first projects spectra into a multi-dimensional feature space of orthogonal latent vectors (Wold et al. 2001) and then links such latent vectors with associated plant traits as dependent variables. Since hyperspectral spectra contain highly correlated bands with redundant information, selecting an optimal number of latent vectors is required to obtain the best model prediction result while avoiding model overfitting (Wang et al. 2020, Gold et al. 2020). In this study, the number of latent vectors was determined through a k-fold cross-validation evaluation of the root mean square errors of prediction (RMSE). A k = 10 was used for both leaf and canopy PLSR models, except for the August 2016 image, of which k = 5 was used due to the limited number of ground training samples (22 samples). The best PLSR models were then applied to the entire image to obtain five trait maps.

### 2.7 Topographic position index

The use of elevation in previous studies does not represent the small-scale topographic variability of the terrain at the same elevation, one of the most important factors that control soil condition and plant biophysical and biochemical processes, especially for shallow-rooted grassland species. In this study, the topographic position index (TPI), a more comprehensive measure of topographic variability proposed by Beven and Kirkby (1979), was used as the means to study the effect of small-scale topography on the trait variation and drought response. TPI, derived from an 8-m DEM, was calculated as 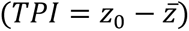, where *Z*_0_ is the difference between a pixel and the center point and 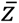 is the predefined neighbourhood’s mean elevation of a radius (Weiss 2001). TPI ranges from-1 to 1, representing different topographic reliefs. Negative values represent lower slope positions, positive values represent upper slope positions, while values near zero represent mid slopes or plateaus (Pierick et al. 2020). The index can be classified into six meaningful categories (Table 1), including valley (VL), toe slope (TS), flat (FL), midslope (MD), upper slope (US), and ridge (RD) that are associated with topographically regulated hydrological processes using the classification scheme proposed by Weiss (2001).

**Table 1.** The classification of topographic position index (TPI).

| TPI category | Abbreviation | Criterion |
| --- | --- | --- |
| Ridge | RD | $z_0 > 1SD$ |
| Upper slope | US | $1SD \geq z_0 > 0.5SD$ |
| Midslope | MS | $0.5SD \geq z_0 \geq -0.5SD$, slope $> 5^\circ$ |
| Flat | FL | $0.5SD \geq z_0 \geq -0.5SD$, slope $\leq 5^\circ$ |
| Toe slope | TS | $-0.5SD > z_0 \geq -1SD$ |
| Valley | VL | $z_0 < -1SD$ |

The correlation between TPI and soil moisture is illustrated in Figure S1. As seen, there was a credible correlation between TPI and soil moisture, with higher soil moisture being associated with lower slope positions and vice versa. Also, the soil moistures of the drought year 2016 were much lower than those of the normal year 2017 but still maintained the pattern.

### 2.8 Statistical Analyses

Trait maps and trait change maps, by subtracting the trait maps of 2016 by the corresponding maps of 2017, were first presented to illustrate trait variation and drought-induced trait change across space. A positive value in the change maps indicates the impact of drought stress, while a negative value indicates better performance of plants. The Pearson’s r correlation analysis was then conducted to evaluate the association, direction, and significance of association between five traits and TPI values at both ground (with ground-and laboratory-measured traits of field plots) and image (with image-derived traits) level to examine topography-driven trait variation. Next, Principal Component Analysis was used to evaluate the association between the first and second components with topographic categories. Boxplots with significant tests were then used to compare trait values and the degree of topographic effect on trait variation within each species and between species over the six topographic categories in both years 2016 and 2017 individually and collectively. Finally, boxplots with a pair-by-pair t-test were used to assess drought-induced trait changes between the two years and how species responded to the stress across six topographic categories.

## 3 RESULTS

### 3.1 Species classification

The RF species classification result along with a 3D map and boundary of topographic categories are shown in Figure 2. The species classification achieved an overall accuracy of 89.6% and a Kappa of 0.870. Among five species, smooth brome (PA = 86.4%, UA = 97.9%) and milkweed (PA = 87.6%, UA = 91.7%) were the most accurately classified species in the area, followed by grape (PA = 83.8%, UA = 92.2%). Orchard grass and goldenrod had a PA = 100%, but UA values were a little lower at 79.1% for orchard grass and 79.8% for goldenrod. Overall, these high classification accuracies were remarkable in species-level classification, especially in this heterogeneous grassland site where species were small, highly fragmented, and had similar colors and textures. The species classification using narrow-band hyperspectral data was more accurate than that reported in a study by Lu and He (2018), which achieved an overall accuracy of 73% using broad-band multispectral data. This is expected as the spectral and textural separability analysis by Dao et al. (2021a) showed that the spectral and textural properties of these species are more distinct in hyperspectral data. It is observed that there was a distinct pattern in species distribution with topographic variability in the site. Smooth brome was the most dominant (37% cover), followed by milkweed (30% cover) and goldenrod (19% cover), while grape and orchard grass were the least dominant with the same percentage cover of just 7%. Orchard grass and goldenrod mainly dominated lowland and flat areas in the centre and the southern regions of the site, while milkweed was found in a range of topographic gradients from lowland and flat to intermediate topographic positions. Grape and smooth brome mainly dominated upper slope positions, only a small proportion of these species was found in lowland and flat regions.

**Figure 2.**
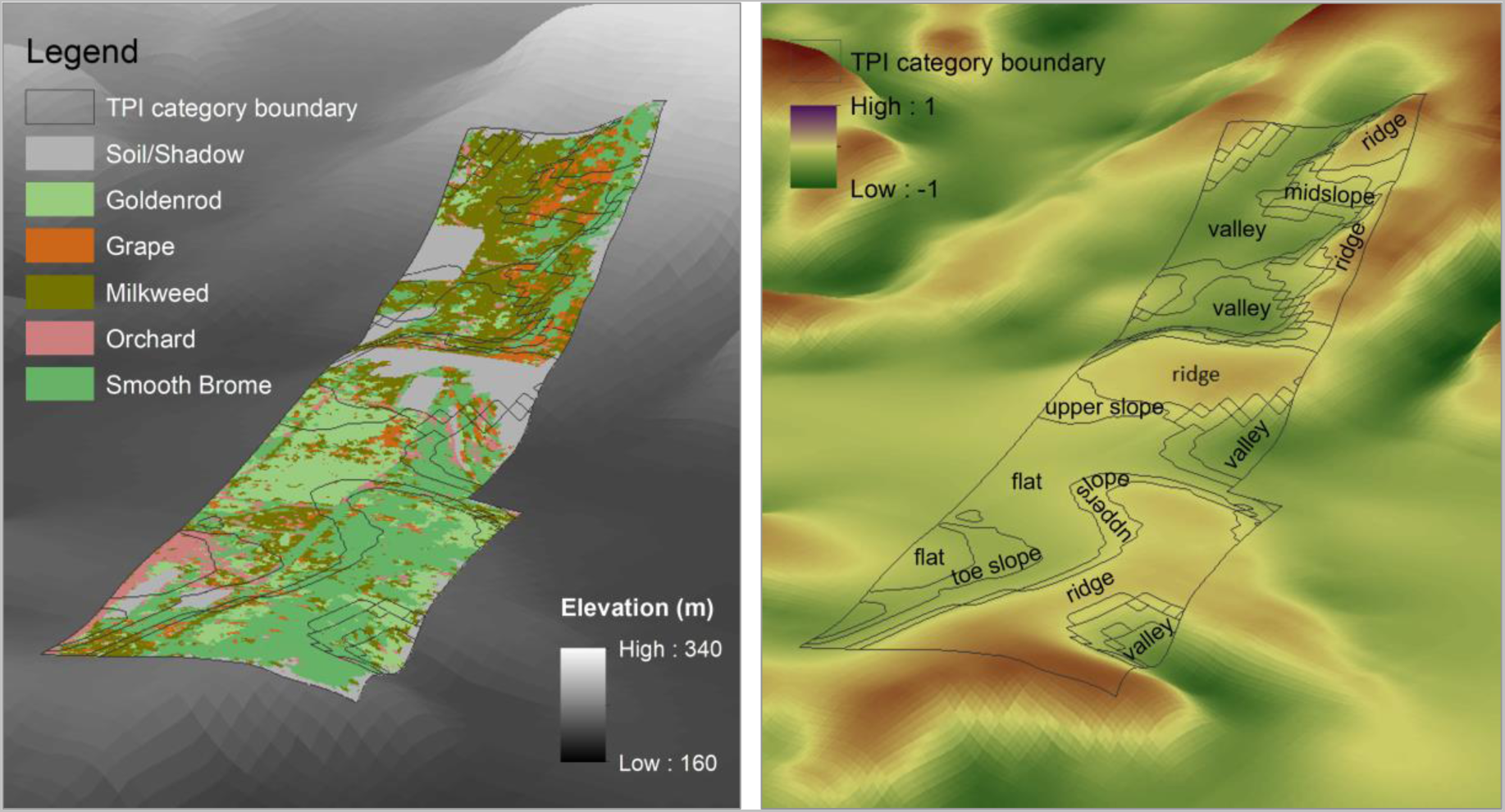
The left figure shows the spatial distribution of native and invasive species overlaid on top of the 3D model of the DEM. The right figure shows the boundaries and labels of six topographic categories on top of the 3D model of TPI.

### 3.2 Plant trait retrieval

Important wavelengths for five traits, detected by PLSR models, are shown in Figure S2. At leaf level (Figure S2a and S2c), wavelengths in visible (∼450 nm and 500-670 nm) and red-edge (700-730 nm) regions were the most important to Chl, Chl/Car, Car, while NIR (750-1000 nm) and SWIR (around 1200 nm, 1430 nm, 1900 nm, and 2400 nm central wavelengths) regions were important to WC. These detected wavelengths were consistent with the results from previous studies (Wang et al. 2020, Asner et al. 2008). Important wavelengths for SLA were found in multiple regions across the spectrum, with some prominent wavelengths at around 750 nm, 1400 nm, 1850 nm, and 2280 nm, which were also reported in previous studies (Wang et al. 2020, Asner et al. 2008). At the canopy level (Figures S2b and S2d), like the pattern at the leaf level, many important wavelengths for all traits were found in the visible (450-620 nm and 650-670 nm) and red-edge (700-750 nm) regions. However, the pattern was different from that at the leaf level in the NIR region (750-1000 nm), where almost all wavelengths were found important to most traits. This relationship in the NIR region was reasonable since vegetation in the area was relatively dense with high LAIs (mean LAI = 2.9 and max LAI = 4.6 in 2016 and mean LAI = 4.3 and max LAI = 7.7 in 2017), causing the dominant control of canopy structure on canopy spectral response (Asner et al. 2008, Croft et al. 2014).

Figure 3 shows the performances of the PLSR model in trait retrieval using spectra at both leaf and canopy levels. At the leaf level, spectral reflectance was able to explain from about 70% to 90.9% of the variation in trait values and the percentage RMSE (%rmse) from 13.8% to 5.5%, except for quantifying SLA of the 2016 data set where 47.1% variation of this trait was explained. Among the five investigated traits, Chl, Car, and WC, with corresponding explained variations of 81.9%, 78.9%, and 88.5%, were most accurately retrieved. Such high retrieval accuracy in these traits was expected as they are among the most abundant and spectrally distinct biochemical properties in plant leaves. At the canopy level, the trait retrieval accuracy decreased to some extent, especially in retrieving Chl/Car of the 2016 data set and Car of the 2017 data set, of which 30% and 47.2% variation, respectively, were explained. The models performed well for Chl with up to 83.8% variation explained and for WC with up to 78.6% variation explained for the 2016 data set. The %rmse varied considerably (between 8% and 30.6%) at the canopy level. The lower accuracy at the canopy level was expected since airborne reflectance was affected by the atmosphere.

**Figure 3.**
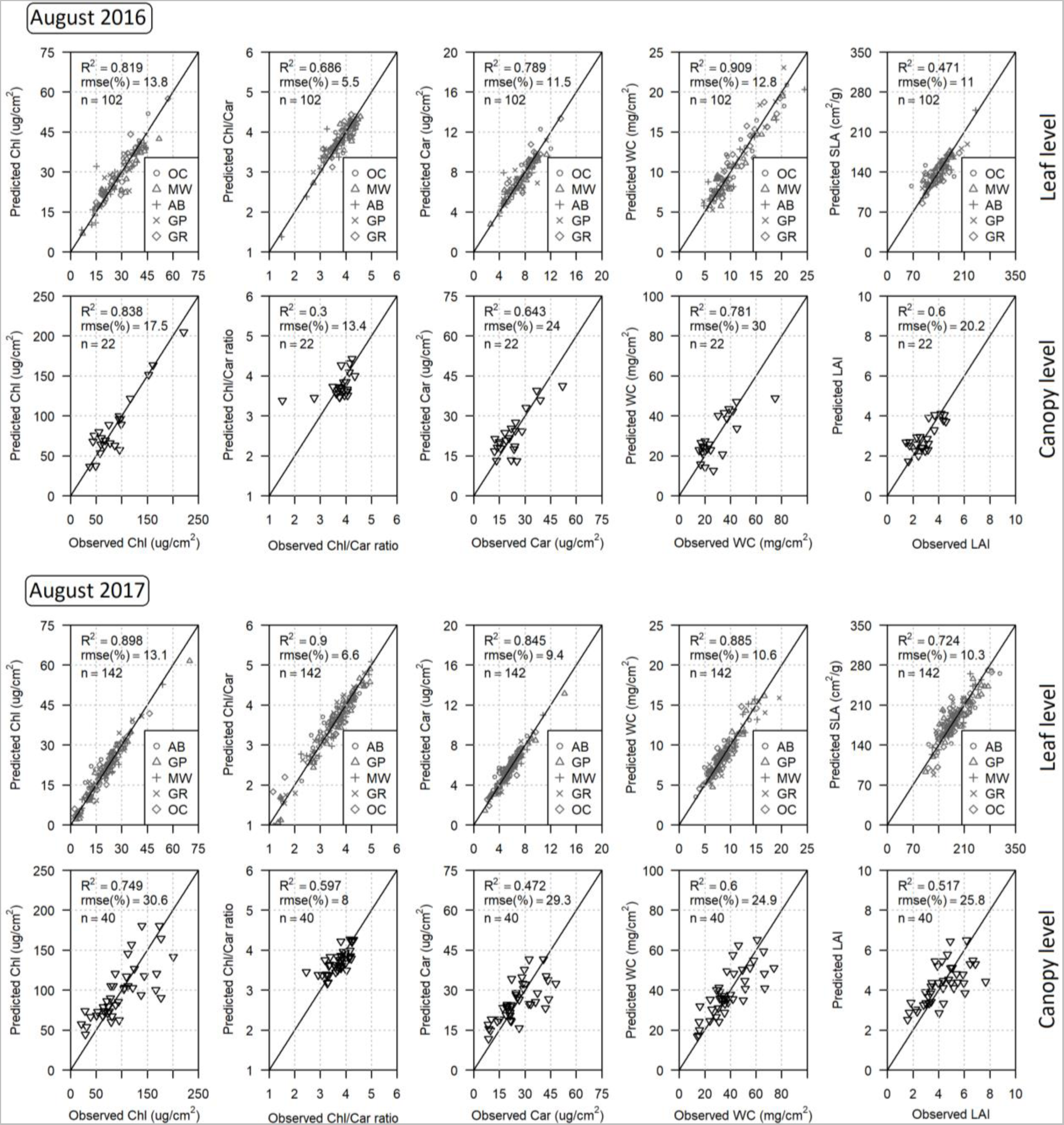
The PLSR results of five traits at leaf and canopy scales for the 2016 and 2017 data. The leaf traits were retrieved by linking leaf reflectance with laboratory trait measurements. The canopy traits were retrieved by linking airborne image-extracted reflectance with calculated canopy traits.

Trait maps, derived from final canopy-level PLSR models, and the change maps between 2016 and 2017 traits are illustrated in Figure 4. Trait values were relatively low in high-elevation or steep-slope regions in the northeast and southwest of the area, where smooth brome and grape were dominant in both years. In contrast, in lowland and flat regions where goldenrod, milkweed, and orchard grass were dominant, especially the central field, trait values were considerably higher. For the change maps, the symmetric color scheme was used where the blue color represents a higher trait value while the red color represents a lower trait value in 2016 than in 2017. In general, trait values in 2016 were much lower than in 2017 over a large proportion of the study area and across topographic gradients, particularly over upper slope positions.

**Figure 4.**
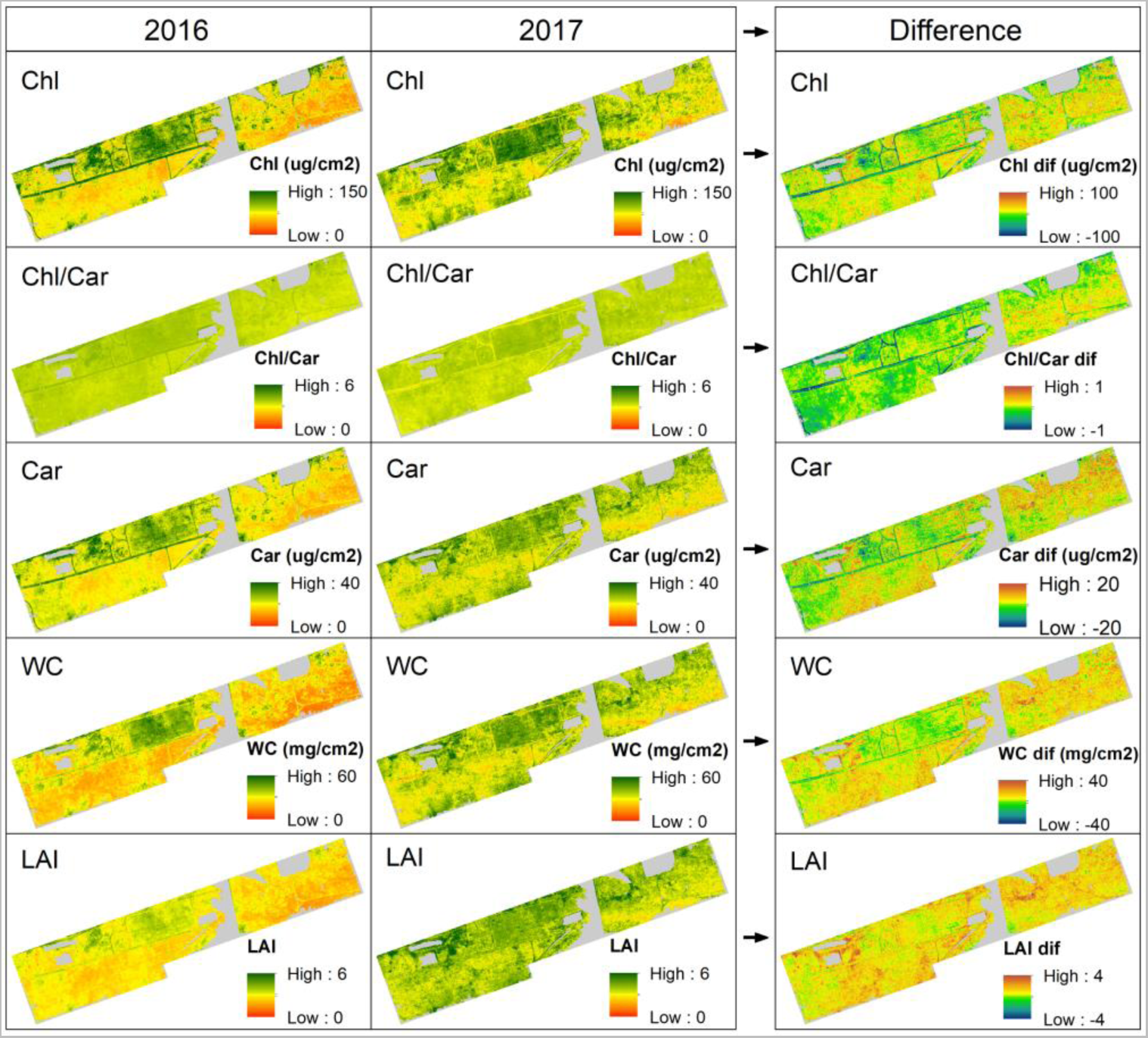
Maps (left and middle subfigures) of Chl, Car, Chl/Car, WC, and LAI and trait difference maps (right subfigures) between 2016 and 2017. The same color scheme and value ranges were used for each map in both years for comparison.

### 3.3 Topographic variability driven trait variation

The plot-level associations between traits and TPI are demonstrated in Figure S3.1. In 2017 (normal year), there were credible negative association between TPI and Chl (r =-0.465), Car (r =-0.495), WC (r =-0.483), and LAI (r =-0.476), while the association was relatively weaker (r =-0.161) in Chl/Car ratio. In contrast, traits of 2016 (drought year) were less affected by topographic variability with negative correlations of r =-0.227,-0.135,-0.112, and-0.208 in Chl, Car, WC, and LAI, respectively. There was almost no correlation between TPI and Chl/Car ratio with r = 0.017, the only positive correlation found in the datasets. Among the five traits, the Chl/Car ratio remained unchanged across topographic gradients in the drought year 2016, indicating the two types of pigments decreased at the same rate. The combined dataset also shows credible correlations between all five traits and TPI.

At the image level, the association between traits, extracted from the derived trait maps, and TPI in five species are shown in Figure 5. Despite some outliers, consistent decreasing trends are observed in trait values from lower slope to upper slope positions in most species. Among those, variations in the traits of orchard grass and goldenrod were affected the most, with a p-value < 0.05 in all traits and all three data sets. Milkweed was the least controlled by topographic variability since correlations were weak and many p-values > 0.05. In particular, invasive smooth brome’s traits were not only the lowest but also remained the most stable over topographic gradients, especially in the drought year 2016 given the regression lines are more horizontal in this species than in others. In grape, trait variation was intermediate in both years. Between the two years, traits of the normal year 2017 were affected by topographic variability at a greater degree than those in the drought year 2016 as the regression lines have higher slopes.

**Figure 5.**
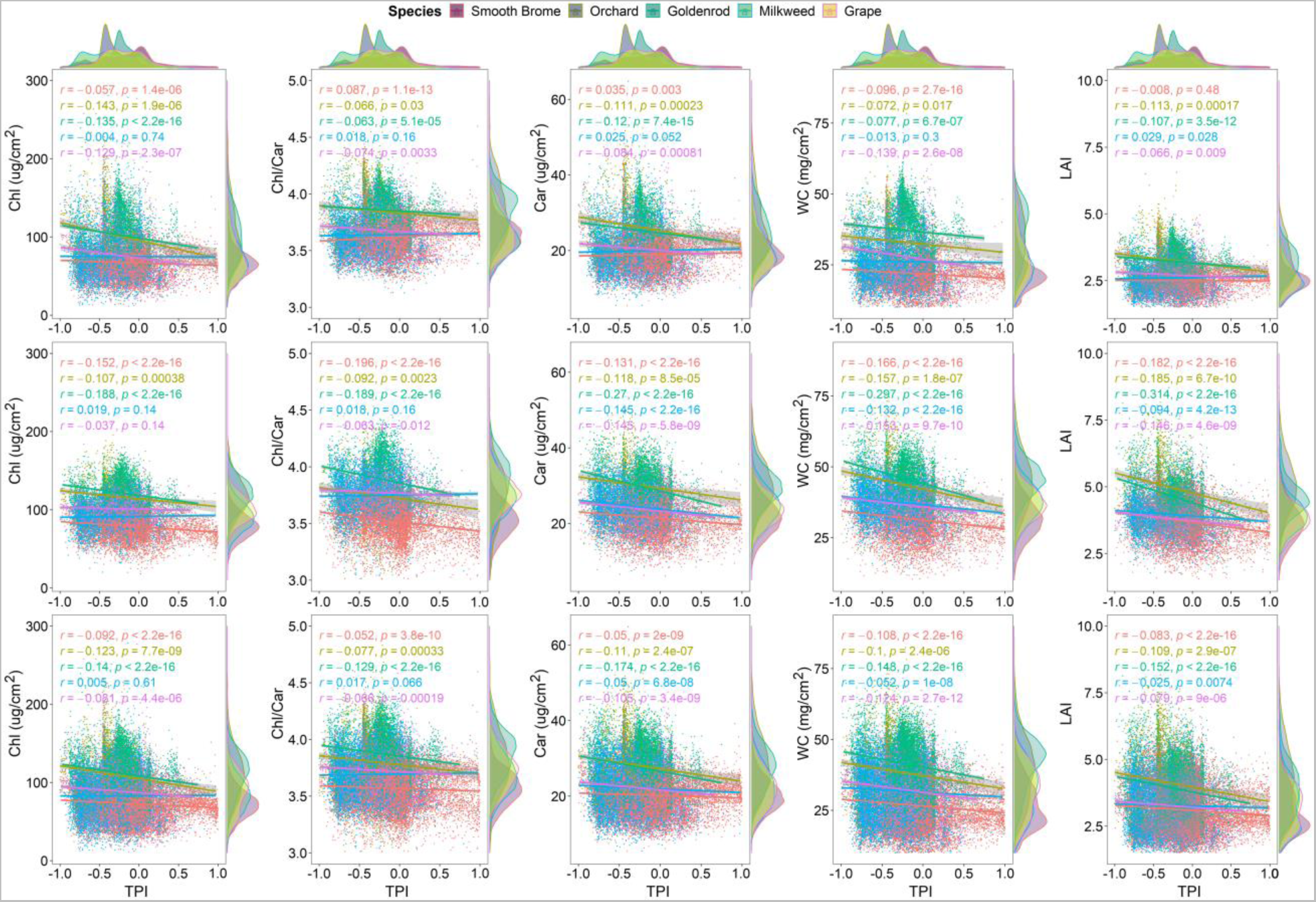
The distribution and association between image-level traits (extracted from trait maps) and TPI, with Pearson’s r correlation and p-value, in five species of the 2016 (top figures), 2017 (middle figures), and combined (bottom figures) data sets.

The principal component analysis of five traits in association with TPI is shown in Figure S3.2. In 2016, Chl, Chl/Car, Car, and WC loaded high on both the first (account for 89.6% of variation) and the second (account for 5.9% of variation) dimension, while LAI was mainly associated with the first dimension. For the 2017 and combined data sets, WC was mainly associated with the first dimension (account for 75.1% for 2017 and 78.6% for combined data set) while other traits loaded high on both the first and the second (account for 19.0% variation for the 2017 and 13.7% variation for the combined data set) dimension. The patterns in all three data sets indicate plants that grew on lower slope positions had higher traits and vice versa.

The within-species trait variation across topographic categories in both the 2016 and 2017 years is illustrated in Figure 6. Overall, there is a clear decreasing trend in all traits from low slope to upper slope positions in 2017. The significance test in Figure S3.4 also shows that most of the traits varied significantly within species across topographic categories, especially between low slope and upper slope positions. In contrast, traits in 2016 seemed to vary largely across topographic regions in almost all species, except in smooth brome. The trend in trait variation from lower to upper slope position was not clear, except in orchard grass, in which trait values at lower slope positions (VL, TS, FL) were considerably higher than at upper slope positions (MS, US, RD). However, the significance test in Figure S3.3 shows that most traits did not vary significantly between three low slope position regions and between three upper slope position regions, but between slow slope and upper slope position groups.

**Figure 6.**
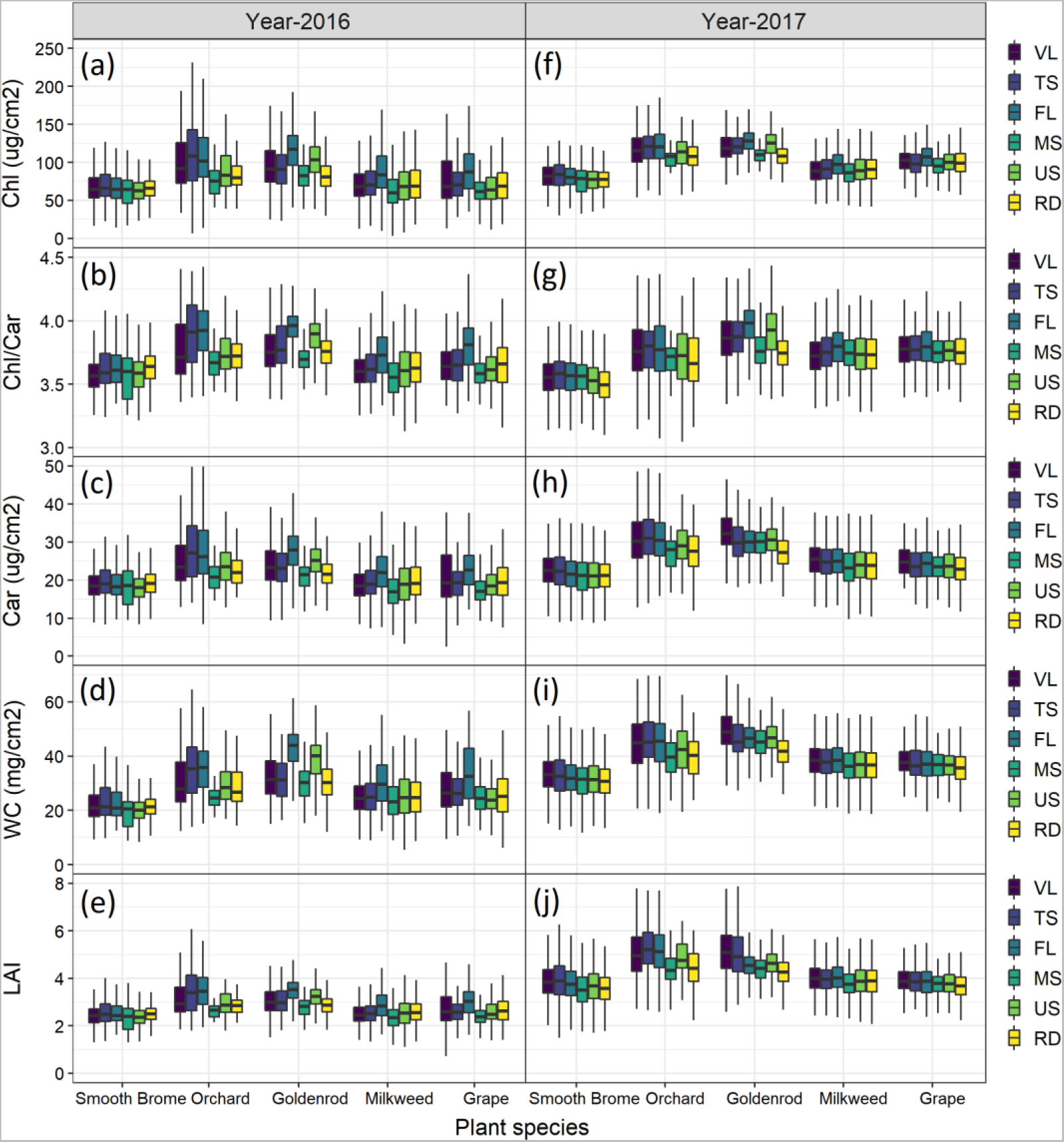
Within-species trait variation in the same year over six topographic categories (VL, TS, FL, MS, US, RD), where (a), (b), (c), (d), (e), and (f), (g), (h), (i), (j) are for Chl, Chl/Car, Car, WC, and LAI of 2016 (left figures) and 2017 (right figures), respectively.

Aside from the within-species variation, topographic variability-driven between-species trait variation is also observed (Figure S3.5). There is a distinct pattern that traits of invasive smooth brome were the lowest, followed by grape, compared to goldenrod, milkweed, and orchard grass in all topographic categories in both years. The significance test in Figures S3.6 and S3.7 also showed the significant differences were found between smooth brome and other species in most topographic regions, especially at upper slope positions, where conditions were relatively dry. Interestingly, smooth brome and grape with lower traits were mainly found at upper slope positions (RD, US, MS), while other species with higher trait values were mainly found at lower slope positions (TS, FL, VL). The test also shows significant differences between most species in most topographic regions.

### 3.4 Drought-induced trait variation

Trait changes, in each species across topographic categories, between the drought year 2016 and the normal year 2017 are demonstrated in Figure 7. Overall, the traits of all species in 2016 were significantly lower than in 2017 in almost all topographic regions, indicating significant impacts of droughts. Exceptions were only observed in Chl/Car in smooth brome (over VL and MS), orchard grass (over VL, MS, US, RD), goldenrod (over RD), and grape (over RD), where the t-test shows insignificant relationships. Out of the five traits, WC and LAI seemed to be affected the most by drought with the highest rate of change, followed by Chl and Car, while Chl/Car was the most stable. There is a distinct trait variation pattern between topographic positions that traits in low slope positions decreased at a greater degree than in upper slope positions in all species. Furthermore, traits of milkweed, grape, and especially smooth brome were considerably lower and remained more stable compared to those of orchard grass and goldenrod, indicating a sign of trait variation within each species and between species under the impact of drought. An example of Chl changes between species in ridges and within smooth brome across six topographic regions in Figure S4.1 also confirms the pattern. As seen, under drought stress, Chl of invasive smooth brome changed less over upper slope positions (RD, US, MS) compared to that over lower slope positions (MS, TS, VL). Also, in ridges the species had lower Chl than those of native species (goldenrod, milkweed, and grape) and drought-sensitive invasive orchard grass in both years; however, under drought impact, its Chl content remained more stable.

**Figure 7.**
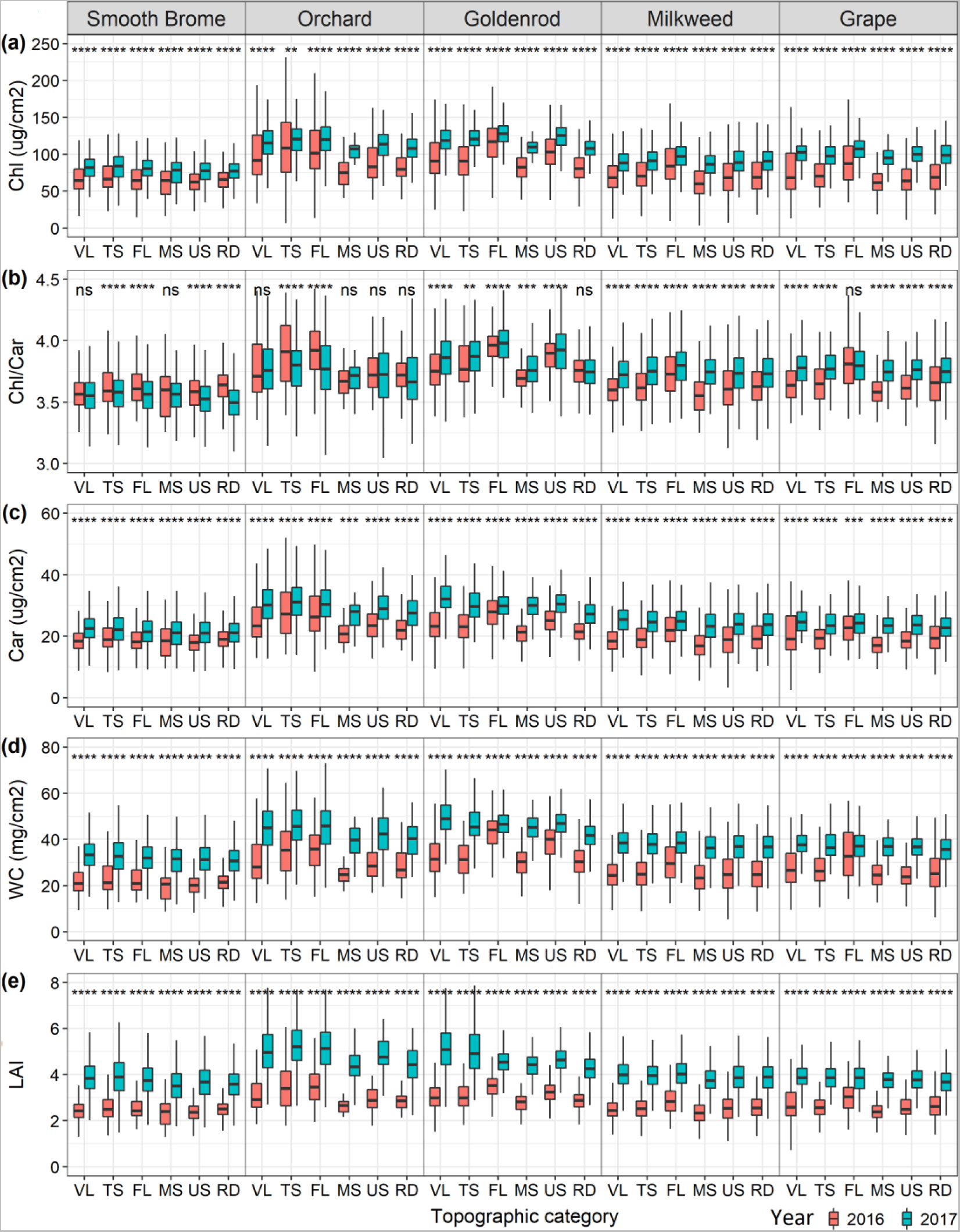
Within-species trait variation between the drought year (2016) and the normal year (2017) across topographic categories (VL, TS, FL, MS, US, RD). Subfigures (a), (b), (c), (d), and (e) are for Chl, Car, Chl/Car, WC, and LAI, respectively.

The comparisons of drought-induced trait change within and between species in each topographic region are demonstrated in Figure 8. Generally, traits changed at different rates among species and over topographic categories. The magnitude and variations of trait changes in lower slope regions were greater than those in upper slope regions. The exception is observed in orchard grass, in which despite experiencing the greatest variations, the changes in this species traits were not significantly different among topographic regions (in significance test in Figure S4.2). Under drought stress, over upper slope regions, WC and LAI changed at the greatest magnitude, followed by Chl, while Car and Chl/Car, experienced less change. Across topographic categories, smooth brome remained more stable, particularly in ridges, under drought stress compared to other species.

**Figure 8.**
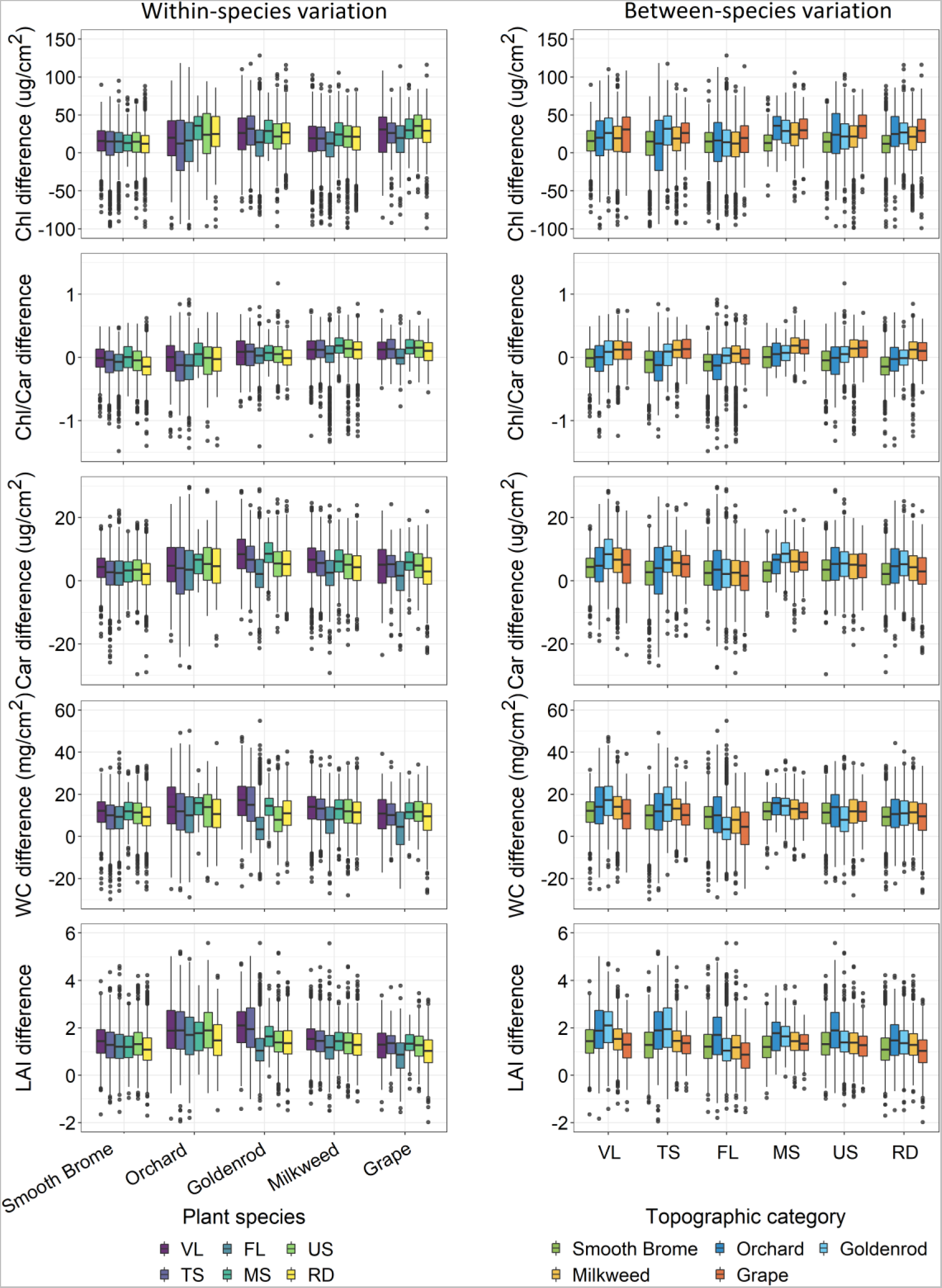
Within-(left) and between-species (right) trait change across topographic categories between 2016 and 2017. From top to bottom are figures of Chl, Chl/Car, Car, WC, and LAI.

Trait changes also varied largely between species in the same topographic regions between the drought and normal year. Generally, traits of smooth brome, which mainly dominated upper slope positions (Figure 2), changed at the lowest rates at almost all topographic positions, particularly at upper slope position regions (MS, US, RD), followed by those of grape and milkweed. Goldenrod that grew mainly at low slope positions experienced the highest rate of change in Chl, Car, WC, and LAI in these regions. The changes in Car, WC, and LAI of orchard grass were among the highest, especially in upper slope positions. Besides, the change in orchard grass varied largely around the median values. The significance test in Figure S4.3 suggests that the changes in invasive smooth brome’s traits were the most distinct that significantly differed from those in native species (goldenrod, grape, and milkweed) in most of the topographic categories. The test shows a few insignificant differences between smooth brome and orchard grass (invasive species) in some categories. The differences in trait change rates among native species were found insignificant in many topographic types.

## 4 DISCUSSION

### 4.1 Trait variation across topographic variability

This study confirms the relationship between traits and topographic variability in earlier studies. Overall, the analyses showed that foliar chemical canopy structural traits monotonically decreased from lower slope to upper slope positions. The large variation within a small geographic region would be driven by the distinctions in microclimatic, soil edaphic, micro-nutritional conditions, and soil water availability over topographic gradients (Pierick et al. 2020, Chadwick and Asner 2016, Swinfield et al. 2020). The decreasing trend of nitrogen availability and soil fertility from lower slope to upper slope position discussed in previous studies (Wolf et al. 2011, Werner and Homeier 2015, Pierick et al. 2020, Swinfield et al. 2020) supports this conclusion. The effect of topographic variability on trait variation was independent of climatic conditions as it is found in both drought year 2016 and normal year 2017. However, the degree of the effect differed between the two years that topographic variability was less influential in 2016 compared to 2017, due primarily to the drought impact. Figure S1 also shows soil moisture in 2017 was higher and varied more largely from low slope to upper slope positions compared to those in 2016.

The effect also varied largely among species. In 2017, smooth brome traits were lower and decreased substantially across topographic positions; however, those in 2016 were mostly topography-independent. This pattern is expected as this invasive species has been reported to be tolerant to drought stress (Sheaffer et al. 1992, Casler et al. 2000). Furthermore, plants that grew at upper slope positions adapted to dry environments, resulting in the stability of traits in these regions. In contrast, orchard grass and goldenrod are less drought-resistant and highly susceptible to water availability (Sheaffer et al. 1992, Werner et al. 1980), resulting in traits being varied to a greater extent, with p-value < 0.05 in the significance test, across topographic positions. The trait variation in milkweed and grape was small in both years. Milkweed mainly grew at lower slope positions with TPIs < 0; therefore, it might not experience substantial variation in soil conditions. Bhowmik and Bandeen (1976) also found the species withstands drought, given its ability to perform well in upper slope regions. Grape has special rootstocks that can adapt to various soil types (Rombough 2002, Ollat et al. 2015). Overall, these analyses indicate that plant tolerance and adaptability are key factors that determine the effect of topographic variability on trait variation.

Previous studies showed an increase in LAI and WC from low to high elevation (Asner et al. 2017, Roderick et al. 2000, Cornwell and Ackerly 2009, Körner et al. 1986), while our study showed the opposite trend. This is because these studies were conducted over various species and ecosystem types at large scales, the impact of small-scale topographic variability on traits could be outweighed by those of soil types, soil fertility, temperature, precipitation, radiation, and humidity (Cordell et al. 1998). In contrast, our study site is relatively small with the same soil type and climatic condition, and trait variation was mainly driven by small-scale topographic variability. Hence, the decreasing trend in LAI and WC could be also due to the impact of prolonged drought and low soil water availability at upper slopes. Plants under these conditions can adjust leaf area and thickness to reduce the radiation load to avoid oxidative damage (Trlica and Biondini 1990) when stomata close to reduce water loss under soil water deficit. The decreasing trend in Chl in our study is consistent with a previous study by Asner et al. (2017) that also used remote sensing to study functional traits. From the field survey, we also observed that leaves at upper slope positions were considerably smaller and thinner than those at lower slope positions. Among five traits, Chl, Car, and LAI seemed to vary the most over topographic gradients, followed by WC. Chl/Car was the most stable, indicating the plant strategy to balance the tradeoff between photosynthesis-driven light absorption and harvesting and photo-oxidative excess light quenching across topography.

### 4.2 Topographic control of drought response

Given the degree of the topography-driven adaptability of species deferred, the response of species is expected to vary when drought occurred. This factor can drive distinctions in trait response within and between species. Figure 7 and Figure 8 showed the underlying within-species distinction in the topographic control of drought-induced trait variation. Accordingly, drought-induced changes in traits of plants at lower slope positions were greater than those in plants that grew at upper slope positions, especially in orchard grass and goldenrod. An example of Chl in smooth brome in Figure S4.1 showed that plants that grew on ridges, upper slopes, and midslopes changed less and remained stable compared to those that grew on flat regions, valleys, and toe slopes under drought impact. An explanation for this pattern is that plants that grow at upper slope positions may develop a strategy to tolerate dry environments, and when drought occurs, plants tend to be more resistant (Liu et al. 2011, Karcher et al. 2008, Volaire and Lelièvre 1997, Pirnajmedin et al. 2015).

In addition to the within-species response, this study also revealed distinctions in drought response between species and between invasive and native species. Invasive smooth brome not only had the lowest trait values but also changed at the lowest rate in all topographic categories, indicating the ability of the species to better withstand drought compared to other species. Such change in smooth brome was expected as this invasive species migrated from a hot environment in Asia and tends to tolerate drought and high temperature (Sheaffer et al. 1992, Casler et al. 2000). This would be the mechanism that hinders the dominance of this invasive species over upper slope regions – occupying 48.2% area in MS, US, and RD compared to only 3.5% of orchard grass, 16.8% of goldenrod, 23.2% of milkweed, and 8.3% of grape. Orchard grass, goldenrod, and milkweed were impacted the most as trait values varied substantially across topographic regions. It is reasonable as orchard grass is less persistent to drought (Sheaffer et al. 1992) while goldenrod seemed to be more adapted to moist soil (rarely found in dry sites) (Werner et al. 1980), and grape is a drought-sensitive species (Padgett-Johnson et al. 2003) that did not perform well once drought occurred. The changes in milkweed’s traits were intermediate. According to Bahmani et al. (2018), milkweed is resistant to water stress and high temperature, especially in more northerly latitudes (Couture et al. 2015). The drought-induced trait changes were relatively similar among native species (goldenrod, milkweed, and grape) because they were more likely found at similar topographic positions (lower slope positions). Whilst, trait changes between these native species and invasive species (orchard grass and smooth brome) were more distinct in many topographic categories. In contrast, distinctions in drought response between the two invasive species were underlying. This is expected since smooth brome is a drought resistant species (Liu et al. 2011, Karcher et al. 2008, Volaire and Lelièvre 1997, Pirnajmedin et al. 2015) while orchard grass is a drought-sensitive plant (Sheaffer et al. 1992). A study by Dong et al. (2014) that showed smooth brome increased root-to-shoot ratio to improve its drought-resistant capacity under drought stress supports our conclusion.

Among traits, WC and LAI were affected at the greatest degrees. A substantial decrease in LAI could be the sign of leaf rolling and wilting to help reduce radiation load (Trlica and Biondini 1990) to avoid thermal and oxidative damages to plant cells and photosynthetic apparatus, while the decrease in WC was likely due to leaf dehydration. The change in Chl was intermediate under stress, especially in low slope positions, leading to the downregulation in photosynthetic activity as the pigment is responsible for the absorption of light for photosynthesis (Tanaka and Tanaka 2006). It is interesting that Chl/Car and Car decreased largely at lower slope positions but remained stable at upper slope positions. The stability of Car, an important accessory antioxidant pigment for photoprotection (Howitt and Pogson 2006, Anderson and Robertson 1960), at upper slope positions indicates better drought tolerance of plants in these regions. At the same time, the stability of Chl/Car indicates the strategy of plants to balance the tradeoff between photosynthetic light absorption and harvesting and photo-oxidative excess light quenching in these regions.

## 5 CONCLUSION

This is the first study that explores the effect of small-scale topographic variability on plant functional trait variation and how species respond to drought in a grassland ecosystem at the species level using airborne imaging spectroscopy. The high-resolution images allow us to understand trait variation and trait response within and between highly mixed co-occurring species, especially between native and invasive species. This study leads to the following major conclusions.

-Reflectance and imaging spectroscopy with PLSR can be used to retrieve plant functional traits with high accuracy. At the leaf level, spectral reflectance was able to explain from around 70% to over 90% of the variation in plant traits with %rmse of less than 14%, while at canopy level, most traits were retrieved with the percentages of variation explained between around 60% and above 80% and %rmse of less than 30%.
-Topographic variability is the main factor that drives within-species and between-species trait variation. The trait-TPI relationship in both drought and normal years (although stronger in the normal year) suggests the topographic effect was independent of the weather.
-The drought-induced trait differences varied within each species and between species across topographic gradients and categories. This pattern indicates the distinction in drought response and tolerance between species, especially between invasive and native species.

The study greatly contributes to the understanding of how topographic variability drives local trait variation and controls drought-induced species response that determines plant biological and ecological processes, performance, and distribution. Moreover, insights about the functional response of invasive species in this study improve our understanding of invasion mechanisms for predicting invasive species habitats, distribution, and spread – essential information for invasive species management. From this unique perspective, the study sets a methodological basis for the advancements of remote sensing-based large-scale assessments of the effect of topography on trait variation and diversity across ecosystems and climatic zones using satellite hyperspectral data such as Hyperspectral Imager Suite (HISUI), Hyperion, Hyperspectral Precursor and Application Mission (PRISMA), EnMAP, and the future Surface Biology and Geology (SBG).

## ACKNOWLEDGMENTS

We acknowledge the financial support from the NSERC Discovery Program (RGPIN-386183) of the Natural Sciences and Engineering Research Council of Canada.

