## Supplementary material for "Imaging spectroscopy reveals topographic variability effects on grassland functional traits and drought responses": ECY23-1241_Supplemental-materials.pdf

### Appendix S1. Topographic position index and soil moisture

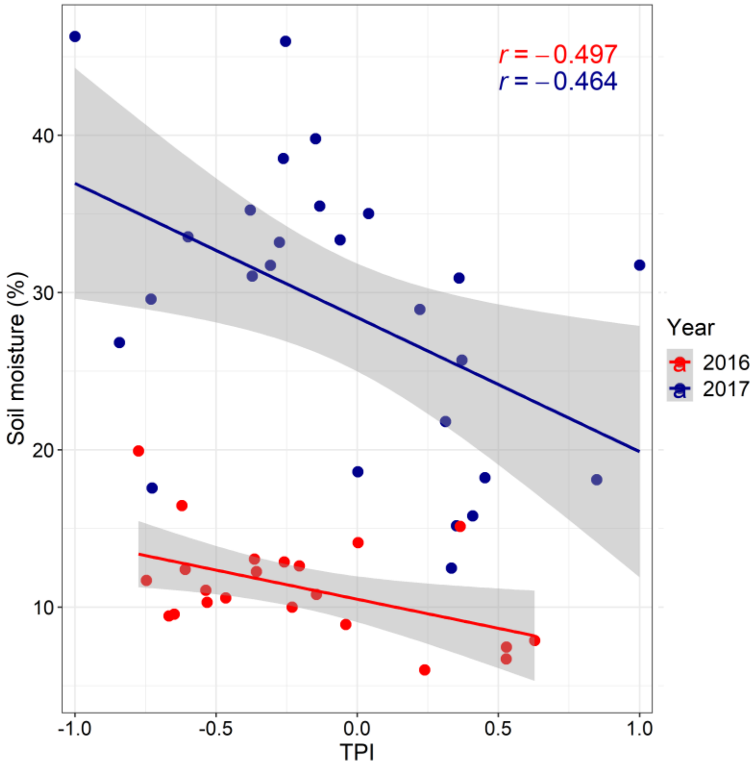


Figure S1. The relationship between TPI and soil moisture at field plots in the drought year 2016 (in red) and the normal year 2017 (in blue). The r values in the figure are the Pearson's correlation between the two variables.

### Appendix S2. Functional trait retrieval from spectral data

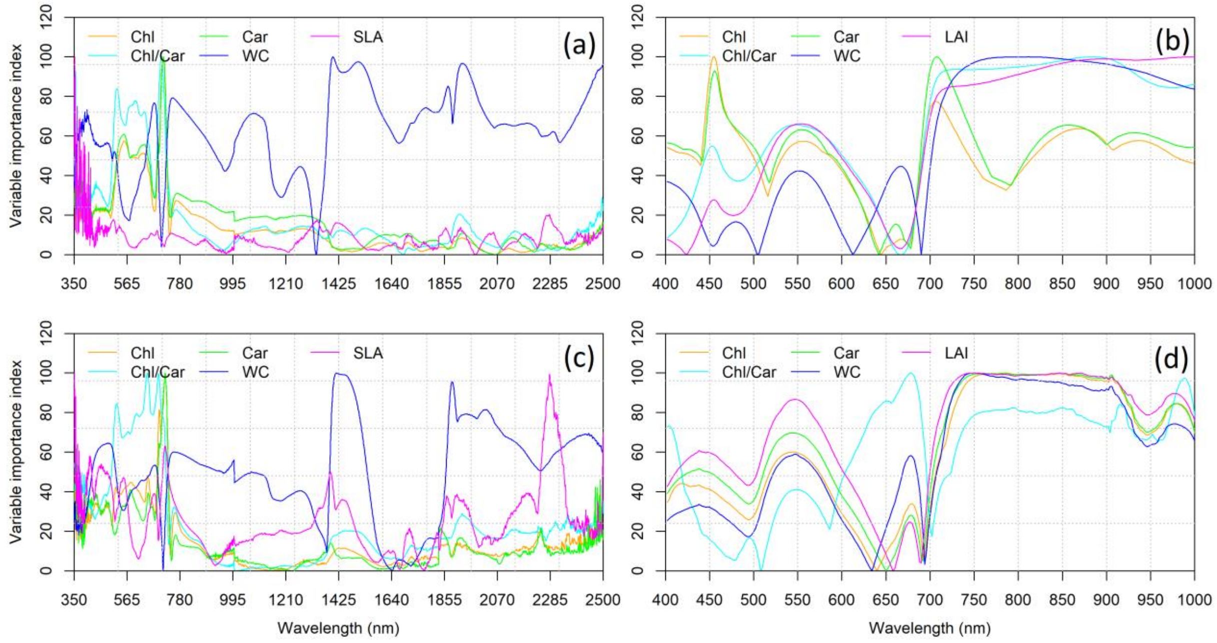


Figure S2. The PLSR variable importance index of each spectral band to five plant traits at the leaf (Chl, Car, Chl/Car, WC, and SLA) and canopy (Chl, Car, Chl/Car, WC, and LAI) levels. (a) and (b) are for the important index at leaf and canopy scales of the 2016 image and (c) and (d) are for the important index at leaf and canopy scales of the 2017 image.

### Appendix S3. Topographic variability-trait relationship

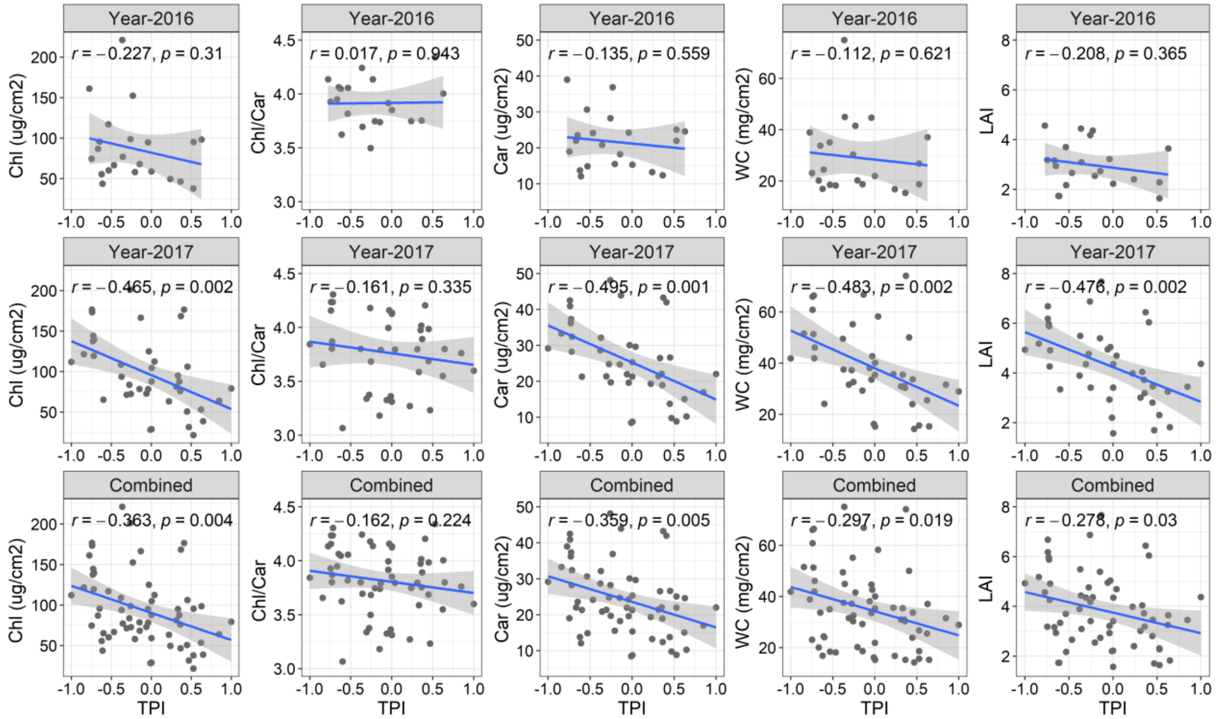


Figure S3.1. The association between five canopy-level functional traits of all five species and the TPI of 2016 (the top figures), 2017 (the middle figures), and combined data sets (the combination of 2016 and 2017 data sets, the bottom figures). The Pearson's r and p-value are also shown in the figures.

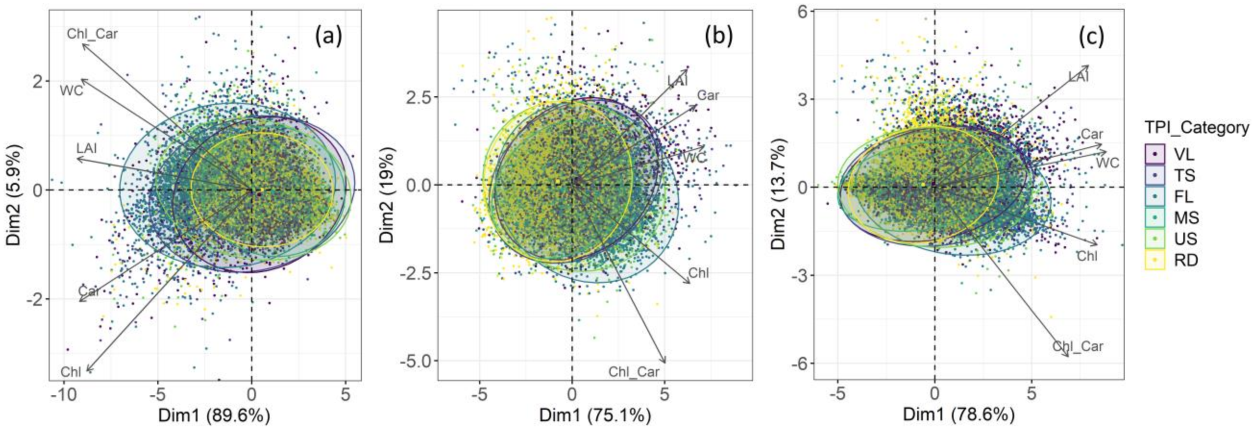


Figure S3.2. Principle component analysis of five functional traits colored by TPI categories. (a), (b), and (c) are for 2016, 2017, and 2016-2017 combined data sets, respectively.

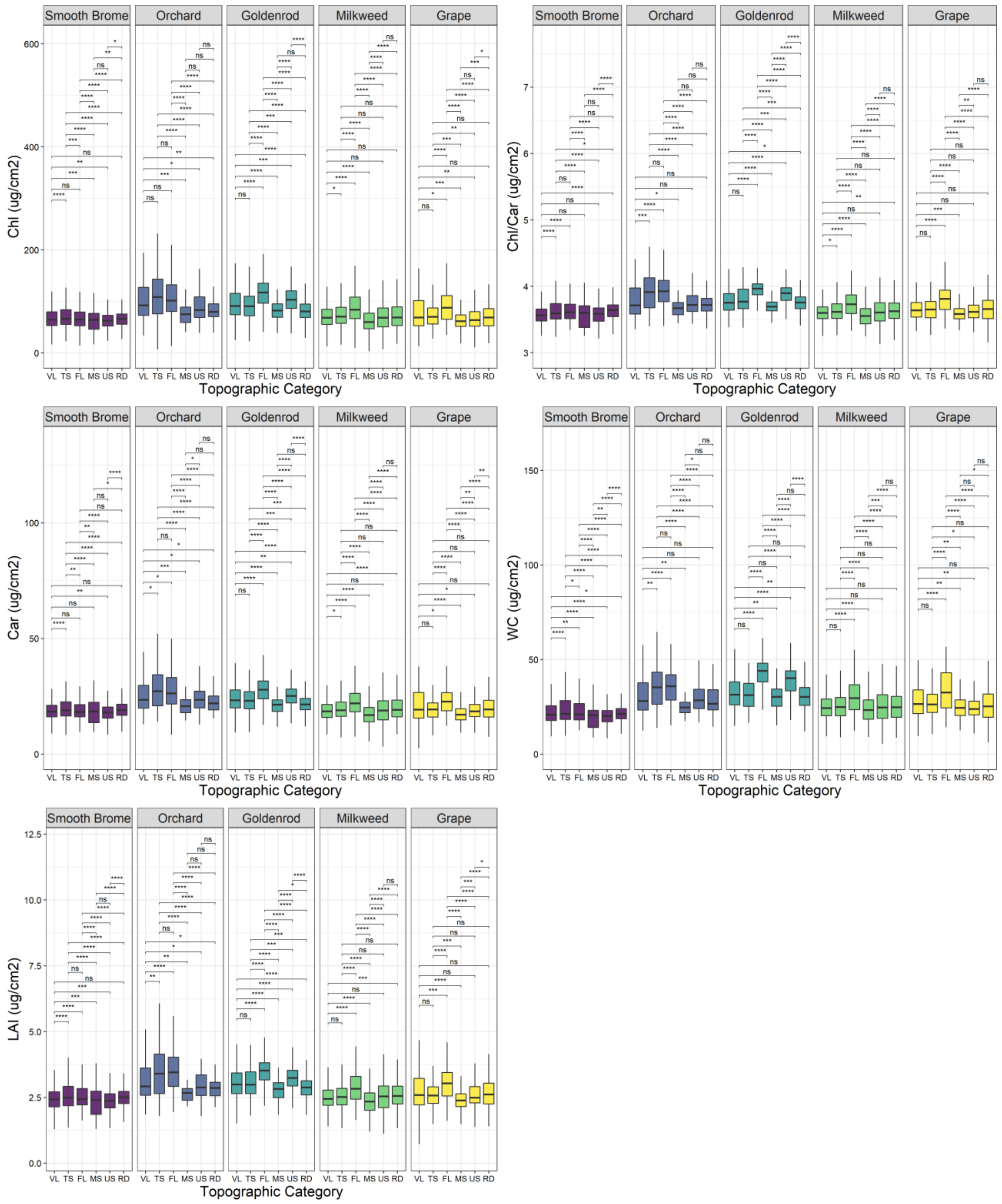


Figure S3.3. The significant test result of the within-species trait variation in five common species across six topographic categories 2016.

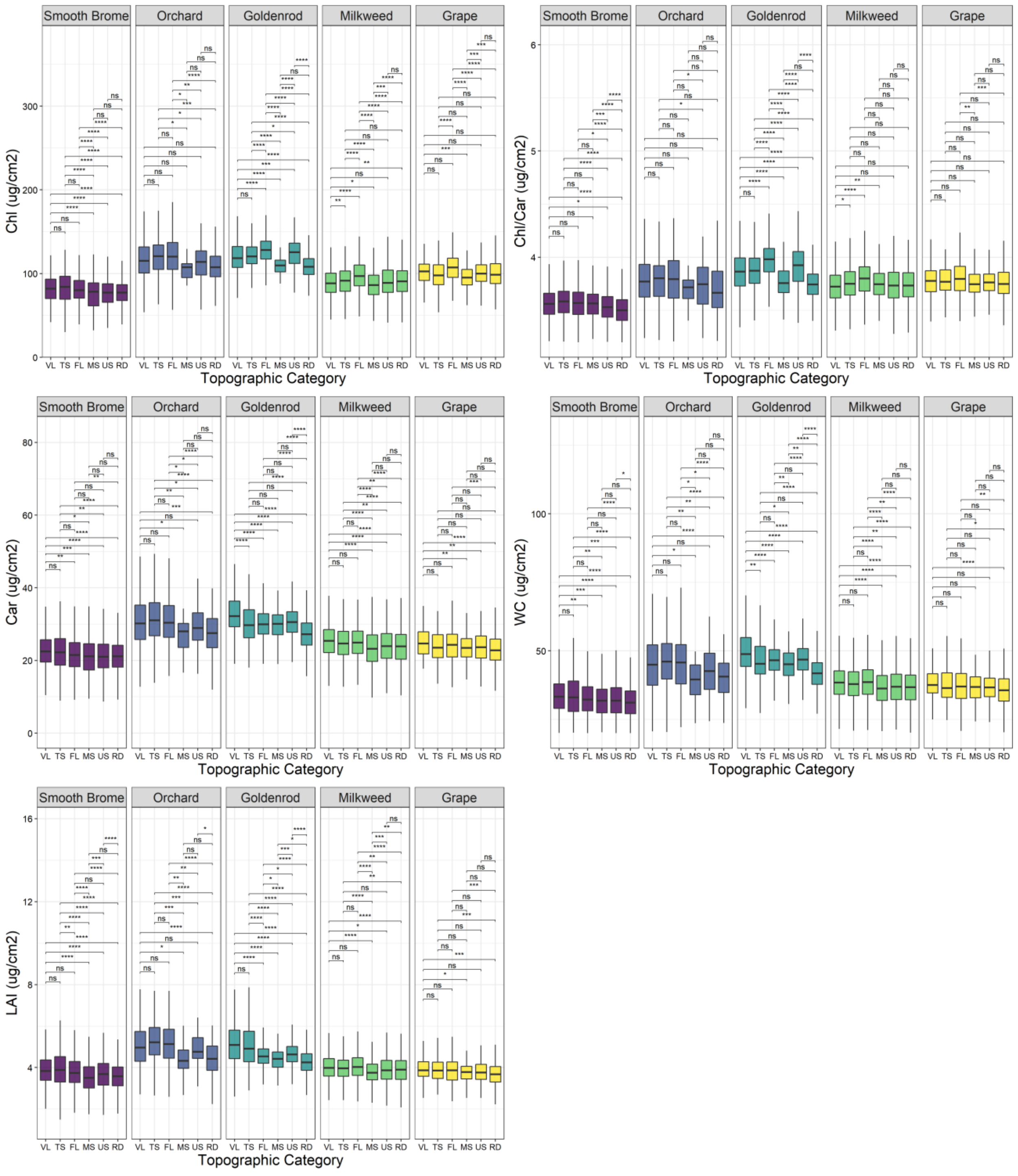


Figure S3.4. The significant test result of the within-species trait variation in five common species across six topographic categories 2017.

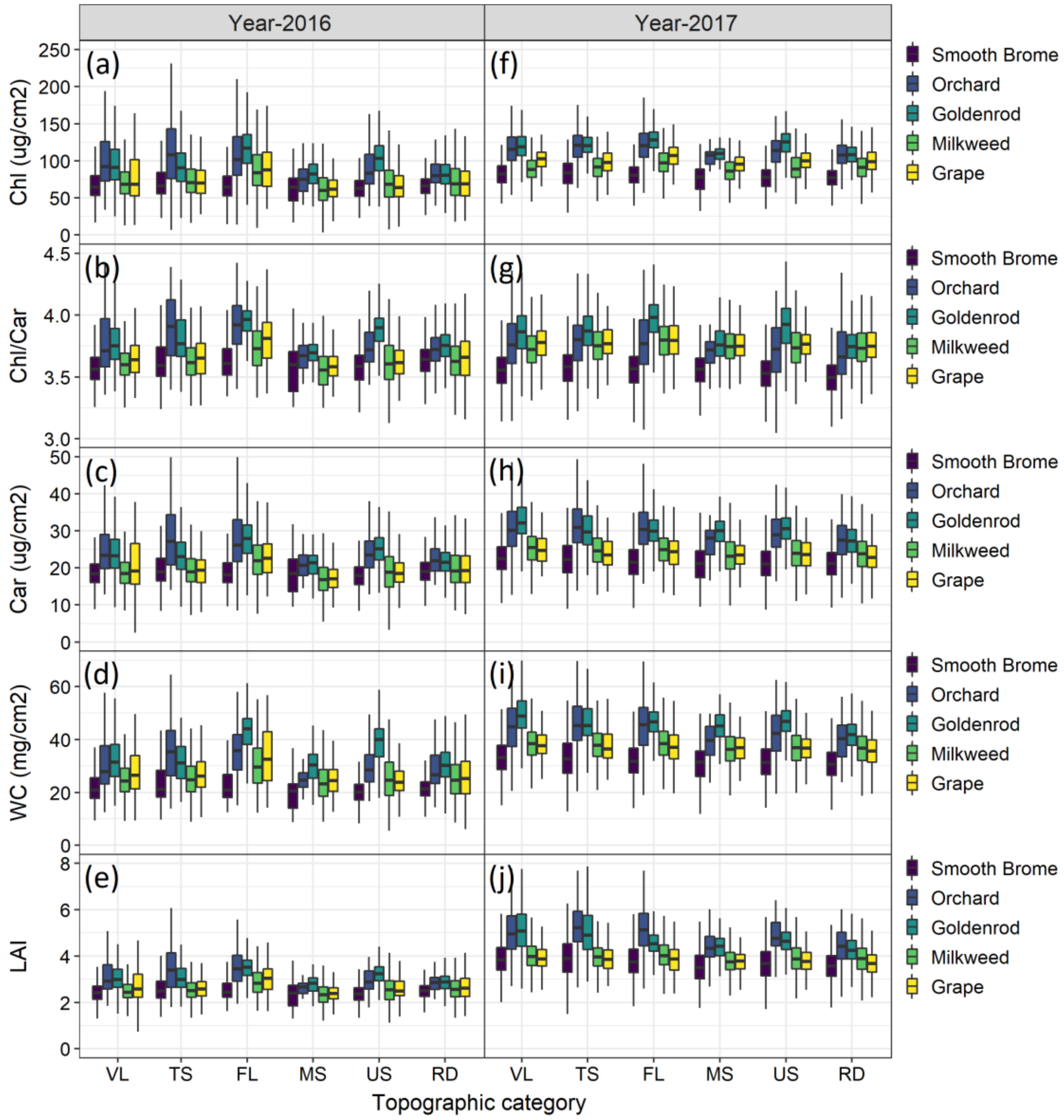


Figure S3.5. Between-species trait variation in the same year over six topographic categories (VL, TS, FL, MS, US, RD), where (a), (b), (c), (d), (e), and (f), (g), (h), (i), (j) are for Chl, Chl/Car, Car, WC, and LAI of the year 2016 (left figures) and 2017 (right figures), respectively.

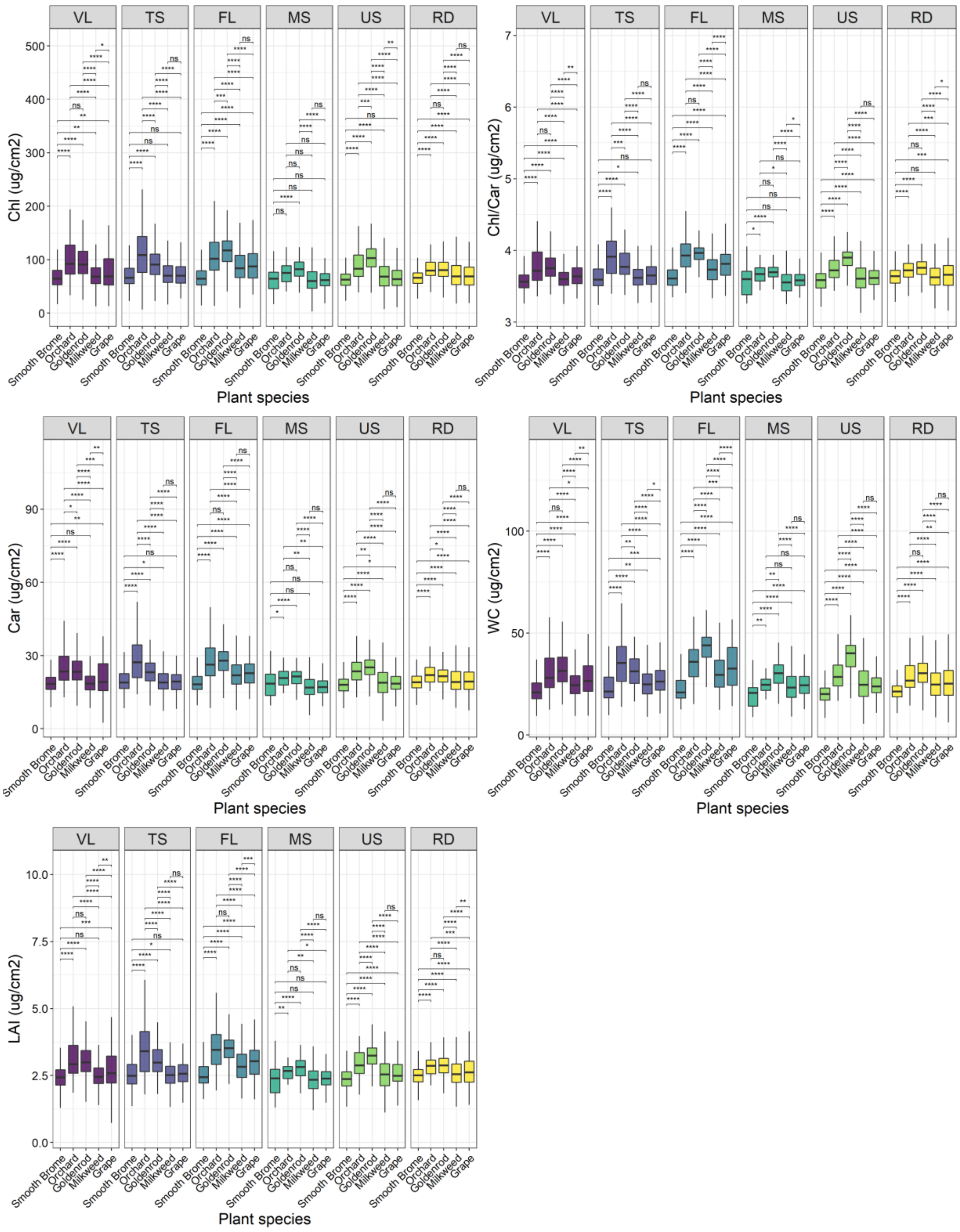


Figure S3.6. The significant test result of the between-species trait variation between five common species in each topographic category in 2016.

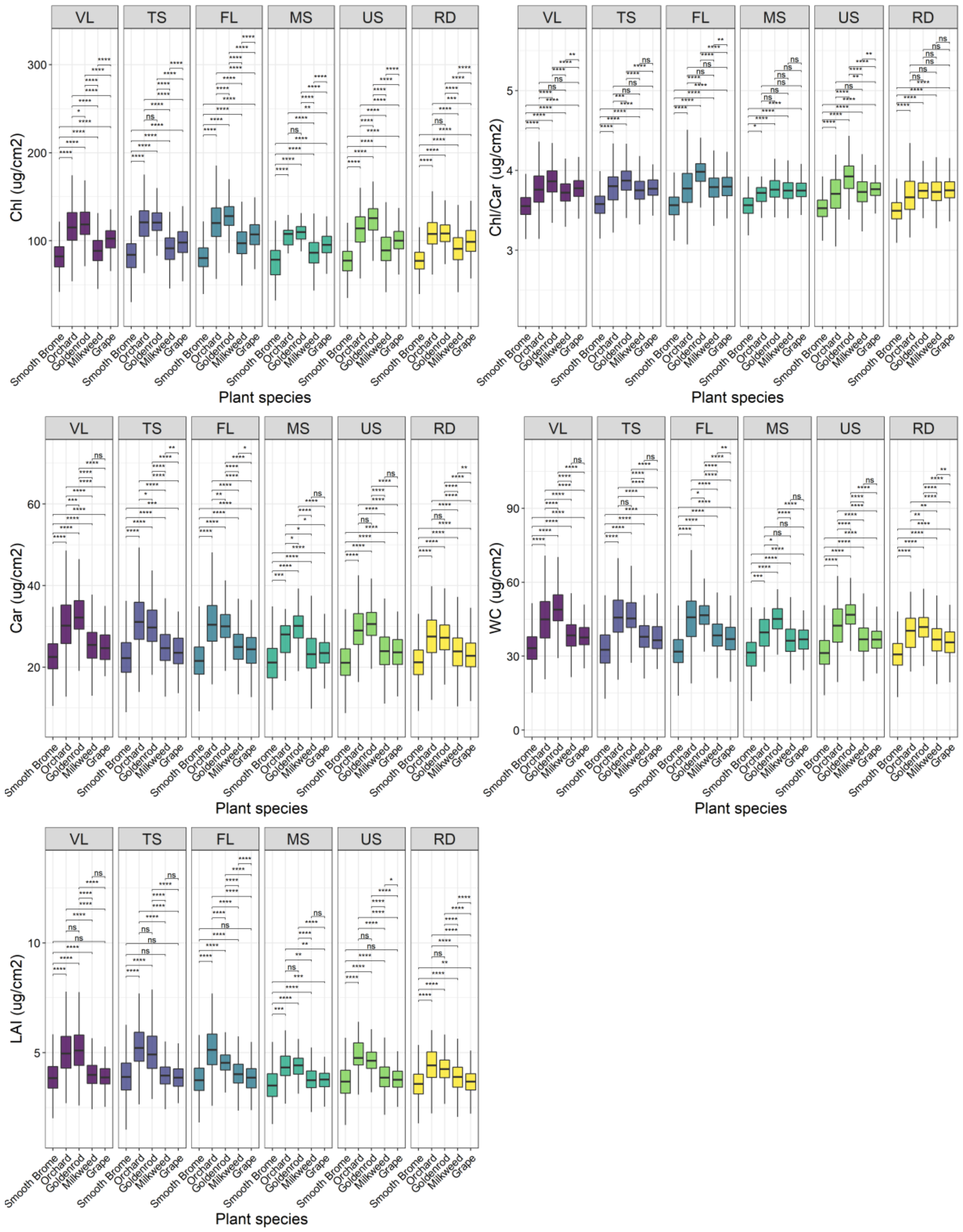


Figure S3.7. The significant test result of the between-species trait variation between five common species in each topographic category in 2017.

### Appendix S4. Drought-induce species response

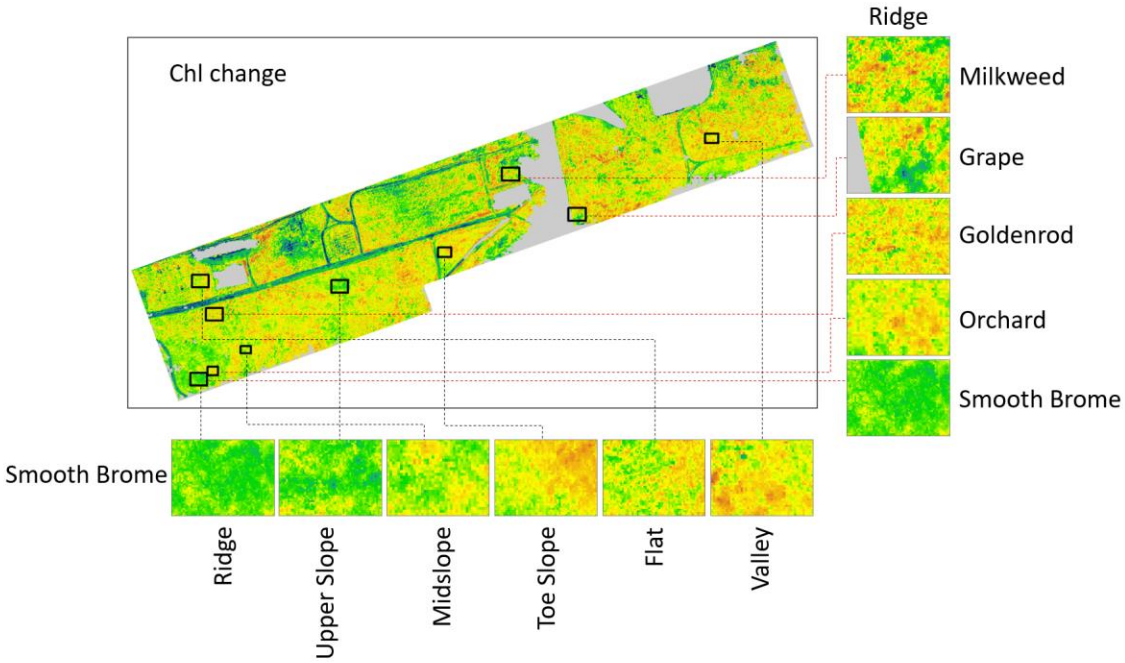


Figure S4.1. An example of changes in Chl between five common species – smooth brome, orchard grass, goldenrod, grape, and milkweed in ridges and within smooth brome between topographic categories (valley, flat, toe slope, midslope, upper slope, and ridges).

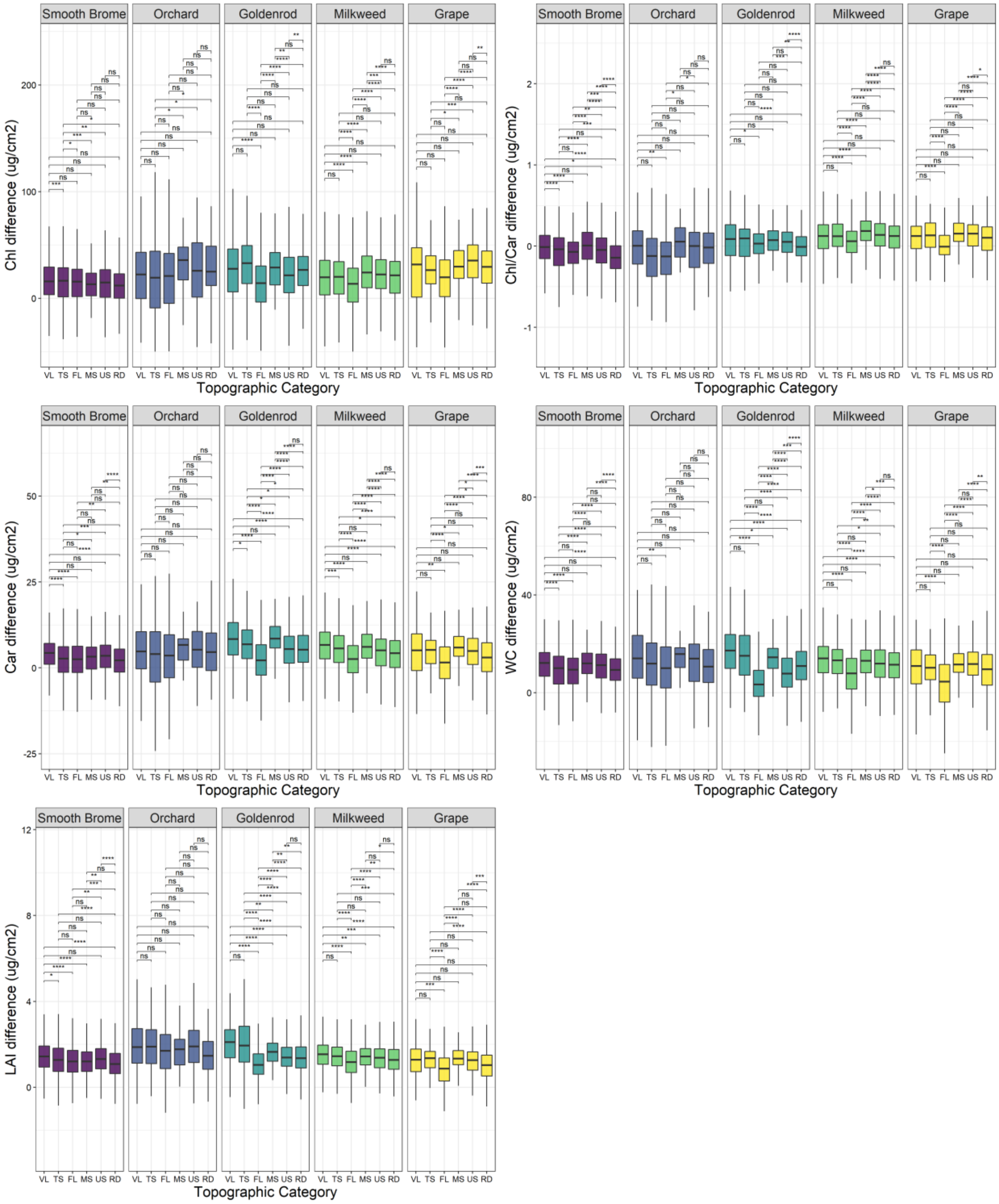


Figure S4.2. Within-species trait change (Chl difference, Car difference, Chl/Car difference, WC difference, and LAI difference) across topographic categories (VL: valley, TS: toe slope, FL: flat, MS: midslope, US: upper slope, RD: ridge) between 2016 and 2017, with the significant tests.

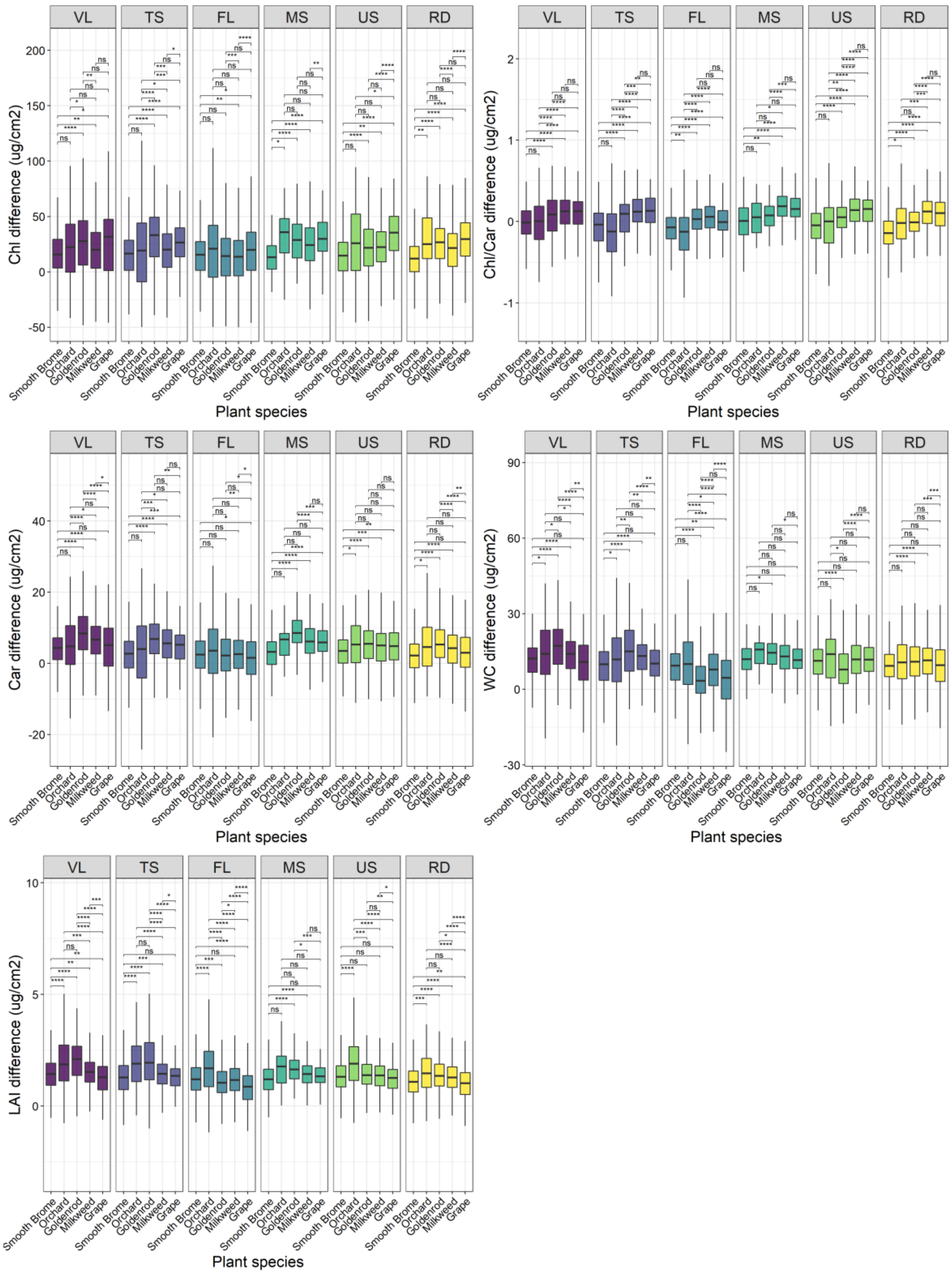


Figure S4.3. Between-species trait change (Chl difference, Car difference, Chl/Car difference, WC difference, and LAI difference) within each topographic category (VL: valley, TS: toe slope, FL: flat, MS: midslope, US: upper slope, RD: ridge) between 2016 and 2017, with the significant tests.
